# Polo-like kinase 1 regulates growth in juvenile *Fasciola hepatica*

**DOI:** 10.1101/2025.07.24.666572

**Authors:** Paul McCusker, Nathan G. Clarke, Rebecca Armstrong, Duncan Wells, Emily Robb, Paul McVeigh, Andreas Krasky, John Harrington, Paul M. Selzer, Nikki J. Marks, Aaron G. Maule

## Abstract

*Fasciola* sp. (liver fluke) are parasitic flatworms that impose a significant burden on the agri-food industry and human health. Immature worms can cause severe damage to the liver as they migrate towards the bile ducts, and yet there is only a single drug to treat this pathogenic life stage, driving the need to identify targets for novel flukicides. Given their crucial role in the growth/development of immature *Fasciola hepatica*, neoblast-like stem cells are an attractive source of new flukicide targets. Kinases are a hugely diverse group of phosphorylating enzymes with key roles in almost all cellular processes. Kinase dysregulation can result in the development of cancerous cells/tissues, linking many to cell cycle-associated proliferation and growth. Here, we annotate the *F. hepatica* kinome, identifying 271 putative protein kinases, representing around 1.6% of predicted *F. hepatica* protein-coding genes with family proportions similar to those of other parasitic flatworms. Many of these kinases, such as polo-like kinase 1 (*fhplk1*), are upregulated in immature worms undergoing rapid growth and development, a process underpinned by the proliferation of neoblast-like stem cells. RNA interference (RNAi)-mediated silencing of *fhplk1* in juvenile liver fluke reduced growth and cell proliferation, suggesting a conserved role within the cell-cycle; the cessation of stem cell proliferation persisted for at least a week following *fhplk1-*RNAi. A PLK inhibitor (BI 2536) was shown to phenocopy the *fhplk1*-RNAi phenotype in a dose-dependent manner, supporting the feasibility of targeting *F. hepatica* neoblast-like cells through kinase inhibitors. Transcriptomic analysis of *fhplk1*-RNAi juveniles revealed 946 downregulated genes, principally associated with the cell cycle or ribosomes. Over 80 of these downregulated genes were also downregulated following juvenile *F. hepatica* irradiation, supporting roles for these kinases in neoblast-like stem cells, and marking them as putative targets for control. Among the 1244 upregulated genes in *fhplk1-*RNAi juvenile worms were many neurotransmitters, receptors and ion channels, exposing the apparent upregulation of diverse inter-cellular signalling systems. While many neurotransmitter pathways promote proliferation in mammalian systems the interaction between neoblast-like stem cells and neuronal signalling in parasitic flatworms remains elusive. Here, the transcriptomic response of *fhplk1-*RNAi juveniles supports a link between neoblast-like stem cell driven growth/development and neuronal signalling.

**Author Summary:** The liver fluke is a parasitic flatworm which causes disease in livestock and humans around the world. While establishing the infection, immature liver fluke migrate through the host liver causing significant damage. Unfortunately, only one drug is currently recommended for treatment of these immature worms, though resistance to this drug is now widespread and exposes the pressing need to develop novel drugs. As worms migrate through the liver, they undergo growth and development which is facilitated by proliferating stem cells. These neoblast-like cells are akin to stem cells seen in other organisms, and as such are responsible for new tissue growth. Many stem cells express kinases, a large family of enzymes that control many cellular processes. In this study we identified all the potential kinases in the liver fluke, including those that may function in neoblast-like stem cells. One of these kinases, polo-like kinase 1 (PLK1), has been linked to cancer development in humans. We used reverse genetics to silence this gene, allowing us to understand its function. Silencing PLK1 reduced worm growth and stopped neoblast-like cell proliferation, confirming that it has an important role in growth and cell division. Also, we showed that a PLK1 inhibitor, developed for cancer treatments, reduced growth and neoblast-like cell activity, illustrating the potential for targeting liver fluke kinases associated with neoblast-like cells with drugs, providing new routes to drug development. We then carried out RNA sequencing of worms after PLK1 was silenced to show the effects of neoblast-like cell loss on the expression of other genes. We found a decrease in the expression of genes that regulate cell division, but an increase in the expression of genes related to inter-cell signalling, including neuronal genes. This supports evidence for interactions between the nervous system and neoblast-like cells which could be exploited in future drug discovery.

## Introduction

Infections with the trematode parasite, *Fasciola hepatica*, can lead to fasciolosis, which is estimated to cause >$3 billion in losses to global livestock yield every year [1]. It also poses a significant threat to human health as a neglected tropical disease [2]. Reports of frontline flukicide (triclabendazole, TCBZ) treatment failure in sheep, cattle and human infections [3], encourage the search for new flukicides, an activity underpinned by novel drug target discovery and validation [4]. This is necessary as TCBZ is uniquely effective in combatting the migrating, highly pathogenic, juvenile parasites which cause the acute stage of the disease. Furthermore, despite intensive research we do not fully understand the mechanisms of TCBZ action or resistance [3,5,6].

Identification of novel targets that disrupt juvenile *F. hepatica* growth/development could underwrite future control programs. Growth in the related free-living flatworms is driven by neoblasts, somatic stem cells that endow species like *Schmidtea mediterranea* with remarkable regenerative capabilities [7]. These neoblasts are sensitive to irradiation [8], and intriguingly, early attempts to develop *F. hepatica* vaccines with irradiated metacercariae restricted development in a mammalian host [9,10]. If this stunted growth in irradiated *F. hepatica* metacercariae was due to disruption of a neoblast-like cell population, it encourages exploration of neoblast-like cells as targets for future flukicides. Over the last decade a range of studies have identified how proliferating stem cells in several parasitic flatworms, including *Schistosoma mansoni*, *Echinococcus multilocularis* and *Hymenolepis diminuta*, facilitate parasite development and tissue renewal/repair, potentially enabling long-term parasitism within hostile host environments [11–19]. Development of an *in vitro* culture platform for juvenile *F. hepatica* enables juveniles to grow/develop towards an adult-like phenotype and showed that *F. hepatica* growth/development was supported by proliferation of neoblast-like stem cells [20]. Furthermore, a recent study demonstrated that irradiation ablates the neoblast-like cell population of *F. hepatica*, resulting in downregulation of stem cell associated transcripts [21].

Numerous protein kinases (PKs) are associated with proliferating cells. PKs are a diverse group of phospho-transferase enzymes that facilitate transfer of the γ-phosphate on an ATP to a substrate’s hydroxyl-group [22]. The human kinome comprises more than 500 PKs [23] that play crucial roles in nearly every aspect of cell biology, with studies suggesting that up to 90% of expressed proteins may be phosphorylated [24]. Functional PKs are typically classed into two groups, the eukaryotic protein kinases (ePKs) and the atypical protein kinases (aPKs) [23]. A staggering 85% of PKs have been linked to cancer or other diseases/developmental disorders [25]. It’s unsurprising therefore that there has been intense development of kinase inhibitors over the last 20 years [26], with over 400 currently in clinical trials (https://www.icoa.fr/pkidb/). The critical role of PKs in many cellular processes, and the abundance of kinase inhibitors make their interrogation as putative anthelmintic targets attractive. Indeed, a protein kinase C-β inhibitor (ruboxistaurin) was shown to kill adult *F. hepatica in vitro* [27]. Kinomes have been assembled for several parasitic flatworms, *S. mansoni*, *Schistosoma haematobium* and *Taenia solium* [28–31], illustrating that all nine major PK families are present, and that the kinome represents a similar proportion of the proteome (1.8-1.9%) to that seen in other eukaryotes [23].

The active role of many PKs in the cell cycle/cell division means that mutations or dysregulation can result in cancerous growths [32]. One set of PKs closely associated with the cell cycle are the polo-like kinases (PLKs), named after a mutant form of the *polo* gene which prevented formation of functional spindle poles during cell division in *Drosophila melanogaster* [33,34]. Additional PLKs have been identified throughout the eukaryotes, from yeasts to humans [35–37], where they are implicated in driving cell-cycle progression [38]. PLKs are serine/threonine kinases with a highly conserved domain architecture, comprising an N-terminal kinase domain and a unique, C-terminal polo-box domain that determines subcellular localisation, and is involved in autoinhibitory regulation of the kinase domain [38]. Some organisms only encode a single PLK (e.g. yeast sp.), whereas others possess up to five PLKs, as in humans [38].

Human PLKs1-3 possess a typical PLK conformation, while PLK4 is more divergent and PLK5 lacks an active kinase domain [38]. Polo-like kinase 1 (PLK1) is most akin to *D. melanogaster polo*, with roles in regulating cell cycle progression from the G2/M transition through to cytokinesis. PLK1 promotes mitosis onset through phosphorylation of substrates that allow downstream activation of the cyclin-dependent kinase 1 (CDK1) – cyclin B complex, driving cell entry into M-phase [29,30]. Other roles for PLK1 include promoting disassembly of the interphase centrosome, formation of mitotic spindle, bipolar attachment of microtubules to sister chromatids, anaphase progression and formation of the contractile ring apparatus for cytokinesis [38,39]. PLK1 has also regularly been observed to be overexpressed in cancer tissues, resulting in its appeal as a putative target for cancer therapies [39].

PLK1 has been linked to cell division in *S. mansoni* and *E. multilocularis* [40–43]. PLK inhibition in adult schistosomes reduced gamete production as well as the viability of adults and schistosomula *in vitro* [40,42]. Inhibiting PLK activity in *E. multilocularis* prevented cell proliferation in mature metacestodes and rendered germinative cells unable to form metacestode vesicles *in vitro* [43]. These investigations suggest that targeting PLK activity could represent a broad-acting anti-parasitic flatworm strategy. A gene displaying typical PLK1 domain architecture has been reported in *F. hepatica* (FhPLK1, [44]) and treatment of *ex vivo* immature and adult *F. hepatica* with the PLK1 inhibitor BI 2536 reduced motility and limited adult egg production [45]. Furthermore, recent transcriptomic studies have linked the *F. hepatica plk1* gene, *fhplk1*, with the neoblast-like proliferative cells of *F. hepatica* as it is upregulated in faster-growing worms [46], and downregulated in worms with ablated neoblast-like cells [21].

Here, we have interrogated available *F. hepatica* genomic datasets to generate a *F. hepatica* kinome, identifying 271 putative PKs. Furthermore, we demonstrate that RNA interference (RNAi) of *fhplk1* undermines the neoblast-like cell driven growth of *in vitro*-cultured juveniles. Transcriptomics of *fhplk1*-RNAi worms highlighted downregulation of many cell cycle effectors, alongside upregulation of inter-scell signalling pathways. Additionally, we show that treating juvenile *F. hepatica* with the PLK1 inhibitor BI 2536 *in vitro* phenocopies the effects of *fhplk1*-RNAi by disrupting neoblast-like cell driven growth. Together, these data support the view that FhPLK1 has a conserved role as a cell cycle regulator in *F. hepatica*, enhancing its drug target candidature as a means of disrupting *F. hepatica* growth/development. Furthermore, our transcriptomic datasets facilitate identification of additional drug target candidates and promote our understanding of interplay between *F. hepatica* neoblast-like cells and other tissues.

## Results and Discussion

### The F. hepatica kinome

Our bioinformatic screen (S1A Fig) of available *F. hepatica* genomes has enabled us to generate a putative *F. hepatica* kinome. In total, 271 PKs were identified (S2 Table; S1 File) with 254 ePKs and 17 aPKs. This constitutes ∼1.6% of the predicted *F. hepatica* proteome, a little smaller than the 1.8-1.9% seen in *H. sapiens*, *S. mansoni*, *S. haematobium* and *T. solium* [23,28–30]. This could be due to the large size of the *F. hepatica* genome, relative to other pathogens [47], or the incomplete nature of the genome assembly, as the total number of PKs identified is comparable to that of *S. haematobium* (269, [30]) and *S. mansoni* (253; updated genome assemblies have removed putative PKs since Grevelding et al. [28]). BLAST searches revealed that 208/253 predicted *S. mansoni* kinase genes had a matching hit in our *F. hepatica* kinome. Members of all nine ePK families were represented, with calcium/calmodulin-dependent protein kinase (CAMK) and CMGC families found to be the largest (Fig1A), representing around 30% of all ePKs. These proportions are similar to those of other parasitic helminths, such as *T. solium, Schistosoma* spp., *B. malayi* and *H. contortus* (S1B Fig; [28–31,62,63]). This contrasts with free-living species (*C. elegans*, *D. melanogaster* and humans), where tyrosine kinases (TK) are the largest family (S1B Fig; [23,48,49]), suggesting a reduced role of TKs in parasitism. We also noted a contrast between the kinomes of ecdysozoans and lophotrochozoans/deuterostomes (S1B Fig), with an expansion of the casein kinase I (CK1s) and receptor guanylate cyclase (RGCs) families in the ecdysozoans, potentially indicating roles for CK1 and RGC in the regulation of moulting.

The largest *F. hepatica* ePK family (CAMK) contains both substrate restricted and multifunctional kinases with a range of homeostatic roles [50]. Due to their critical roles in Ca^2+^ signalling pathways, they have been proposed as potential anthelmintic targets. Schistosome CAMKII increases its expression/activity following treatment with the paralysis-inducing drug praziquantel, with RNAi reducing motility/viability [51,52], suggesting that CAMKII plays a role in worm motor function/motility. The CMGC family is the second largest ePK family in *F. hepatica.* This highly conserved family consists of members that play critical roles in cell cycle control, such as the cyclin-dependent kinases (CDKs), mitogen-activated protein kinases (MAP kinases) and CDK-like kinases (CDKLs). This high degree of conservation is unsurprising given the crucial role these kinases play in fundamental cell biology, especially in proliferating cells. This may provide opportunities for the development of flukicides that disrupt growth via CDK and MAP kinase inhibition. Indeed, inhibitors targeting these kinases have already been developed for cancer treatment [53], and repurposing/redeveloping them to target the neoblast-like cells of *F. hepatica* and other parasitic flatworms could lead to their use as novel anthelmintics.

We examined the phylogeny of *F. hepatica* ePKs using CLANs software and found the ‘Other’ grouping to be the most divergent (Fig 1A). This was not surprising given the diversity of this group which includes the polo-like kinases, NimA-related kinases and aurora kinases among others. In contrast to this, the CMGC family are the most closely clustered, a reflection of their conserved roles. The AGC, CAMK, CK1 and TK families clustered closely with only a few divergent members. However, clustering was more diffuse for the tyrosine-like kinases (TKLs) (Fig 1A), with a bone-morphogenic protein kinase (FhHiC23_g1262) and an integrin-linked protein kinase (FhHiC23_g8606) notable outliers. These divergent sequences may be druggable targets, especially with only a 19-22% sequence identity when compared to their human orthologues. Recent publications in *F. hepatica* have also demonstrated the potential of targeting kinases (protein kinase C β and p21-activated kinase 4) to disrupt motility/viability [27,54].

**Fig 1.**
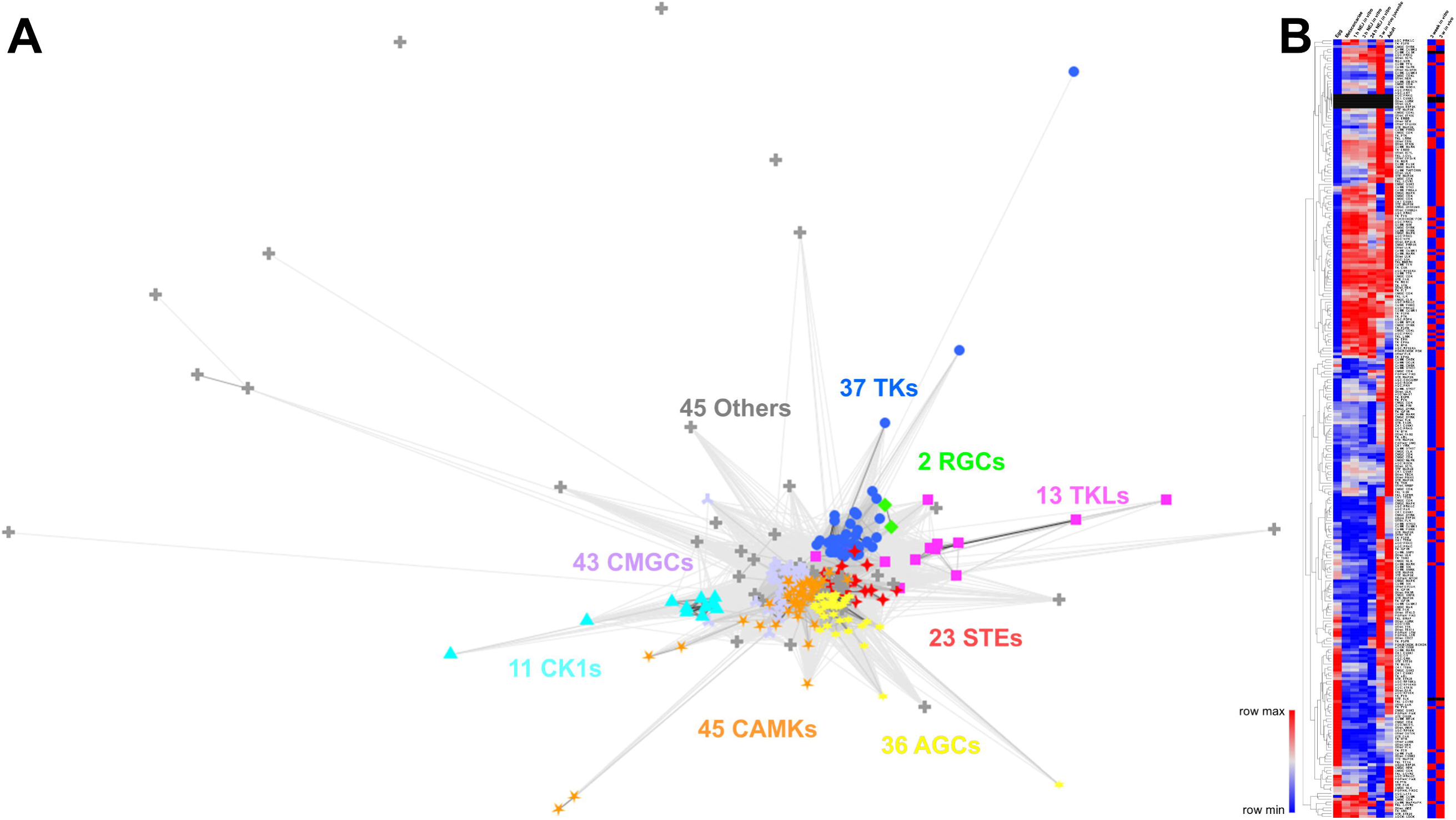
*Fasciola hepatica* kinases are enriched in developing life stages. (A) CLANs analysis (1M iterations) of *F. hepatica* eukaryotic protein kinases shows presence of all nine major families with conservation among CMGC kinases and mitogen-activated protein kinases (STEs) but divergence among tyrosine kinase-like kinases (TKLs), casein kinases (CKs) and calcium/calmodulin kinases (CAMKs). (B) Heatmap of *F. hepatica* kinases (where data was available) in various life stages (Left) with average euclidean distancing applied shows upregulation of many kinases in newly excysted juveniles (NEJs) and immature worms as well as upregulation of the majority of kinases in *in vivo* juveniles compared to *in vitro* juveniles (Right), colours correspond to z scores (red = upregulated; blue = downregulated; grey = no change; black = no expression data).

We also identified 17 putative *F. hepatica* aPKs (S2 Table), functional PKs with a typical catalytic kinase fold that lack sequence similarity to ePKs [55]. As in other organisms, there are considerably fewer aPKs than ePKs, though the number identified here is comparable to that found in *C. elegans* [56]. Several conserved *F. hepatica* aPKs (S1C Fig) play vital roles in cell proliferation, linking them to their neoblast-like stem cells, such as mammalian target of rapamycin (mTOR), ataxia–telangiectasia mutated kinase (ATM), ataxia–telangiectasia and Rad3 related kinase (ATR) and the phosphoinositide 3-kinases [57–59], again highlighting numerous pathways that could be exploited for juvenile fluke control.

### FhPLK expression linked to developing juveniles

We used published life-stage transcriptomic data [47] to examine expression profiles of putative PKs. Of the 266 kinases for which we have life stage transcriptomic data, 116 show the highest relative expression in NEJs or immature worms (migrating three-week-juveniles) (Fig 1B). Seven of these genes with enriched expression are *cdk*s, crucial cell cycle regulators [60]. Among them is a putative *cdk1* gene (FhHiC23_g16347) which has been implicated in regulating the pluripotency of embryonic stem cells [61], and can drive the cell cycle in early embryonic cells lacking all other CDKs [62]. Additionally, four putative *F. hepatica* fibroblast growth-factor receptors (*fgfrs*) and two *plks* are most highly expressed in these developing life stages. We also utilised a published transcriptomic dataset that compared *in vitro F. hepatica* juveniles with faster-growing *in vivo* juveniles [46] and found 81% of kinases were more highly expressed in the faster-growing *in vivo* juveniles (Fig 1B), again supporting the hypothesis that kinases play an integral role in growth/development.

Development-associated kinases were further interrogated using the *F. hepatica in vitro*/*in* vivo transcriptomic datasets [46] alongside a dataset which identified genes associated with *F. hepatica* neoblast-like cells [21]. Six kinases were downregulated following neoblast-like stem cell ablation, and 21 kinases were upregulated in faster-growing worms (S2 Table). Notably, two genes were prominent in both of these datasets, a *fgfr* (FhHiC23_g5821) and *plk1* (FhHiC23_g6132). FGFRs are tyrosine-kinase receptors linked to proliferation [63] with drug inhibition, RNAi or CRISPR-interference in *S. mansoni* and *E. multilocularis* reducing neoblast-like cell proliferation [11,12,64,65]. Three FGFR inhibitors have been approved by the FDA for treatment of cancers and idiopathic pulmonary fibrosis [66], highlighting the druggability of these kinases.

PLK1 is a core cell cycle effector, regulating centrosome assembly, mitotic entry, the spindle, cytokinesis and more besides [39]. PLK1 has been linked to cell differentiation and survival in other parasitic flatworms [40–43], while in *F. hepatica* its downregulation following irradiation and upregulation in faster growing worms (S2 Table) support its link with neoblast-like stem cells. Our HMM screen and additional BLAST searches identified two further putative *F. hepatica* PLKs (FhPLK2 [FhHiC23_g13468] and FhPLK4 [FhHiC23_g6087]; Fig 2A). However, only FhPLK1 was predicted (full sequence confirmed using transcriptomic reads) to possess all signature PLK1 domains (Fig 2C), including the ATP-binding domain [67], catalytic domains, Mg^2+^ chelating domain [68], phosphorylation site [69] and phosphopeptide binding sites [70,71]. Published life stage transcriptomic data [47] show that *fhplk1* expression peaks in eggs (Fig 2B), potentially indicating significant proliferative cell activity as the embryo develops into the unhatched miracidia, like that seen in *S. mansoni* [72]. Following eggs, relative expression was highest in immature worms and adults, life stages with significant neoblast-like stem cell and germ cell activities respectively [46,73,74]. Moreover, transcriptomic data from Robb et al. (Fig 2B, [46]) show greater *fhplk1* and *fhplk4* expression in faster-growing *in vivo* worms, correlating with observed increased cell proliferation. Houhou et al. [44] also found higher *fhplk1* expression in immature worms and adults compared to NEJs (24 h post-excystment) via PCR, though they also observed ∼6-fold increase in expression in adults compared to immature worms. The difference seen here between transcriptomic and qPCR methodologies could be due to low replicate numbers in the published transcriptomic datasets [47]. As *plk1* has previously been localised to *S. mansoni* reproductive structures [40], the increased expression in adult *F. hepatica* is likely associated with germ cell activity in this life stage. Together, these transcriptomic data indicate that *fhplk1* is associated with cell proliferation in *F. hepatica*.

**Fig 2.**
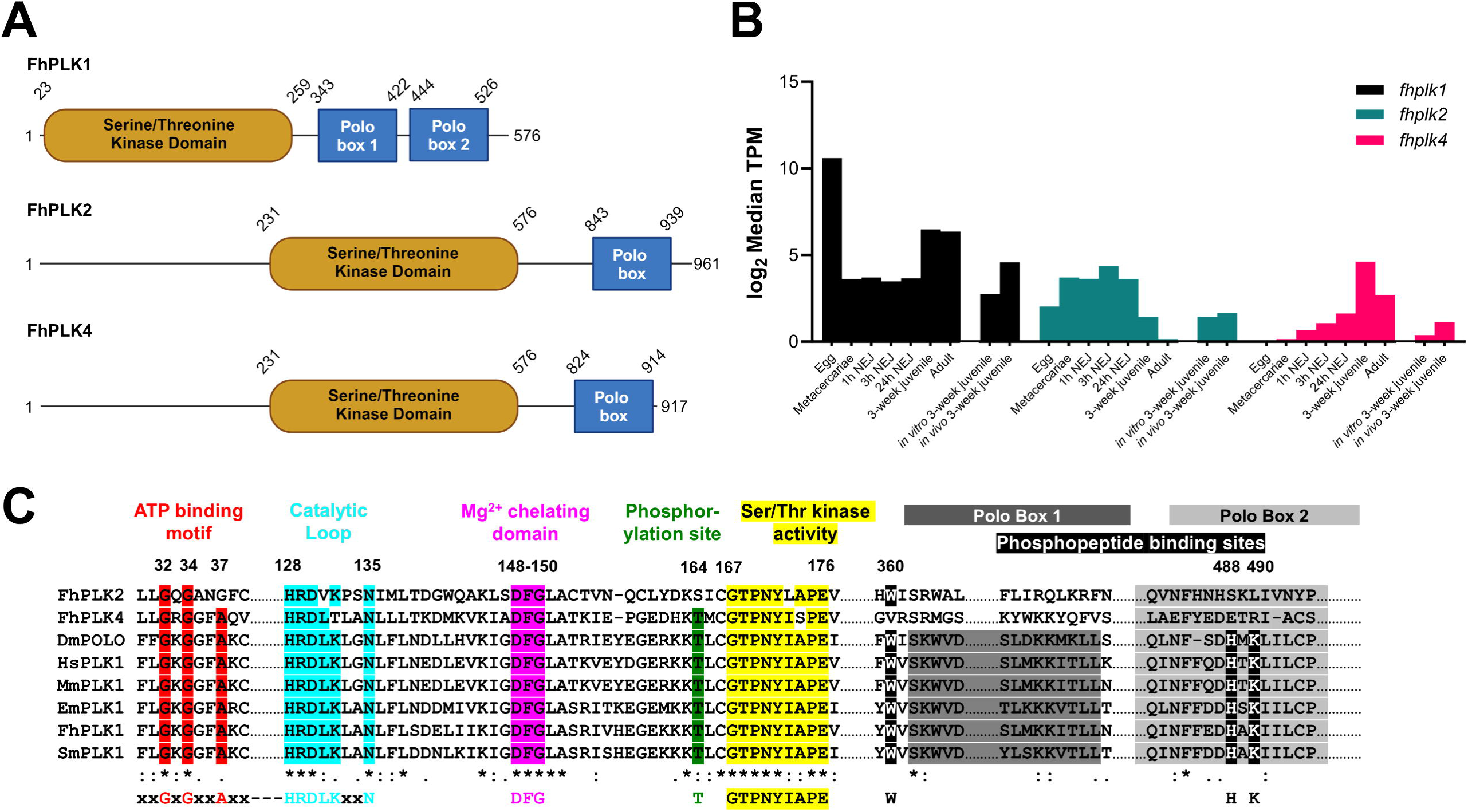
*F. hepatica* polo-like kinase 1 (PLK1) possesses all functional kinase domains. (A) Schematic of *F. hepatica* PLKs, with domains as predicted by Interpro, illustrating N-terminal serine/threonine kinase domains and C-terminal polo-boxes. (B) Expression of *fhplk* genes from published datasets; Log_2_-transformed median transcripts per mapped million reads (TPMs) of *F*. *hepatica plk* genes across various life stages (egg, metacercariae, 1h NEJ, 3h NEJ, 24h NEJ, 3-week juvenile, adult; [47]); Log_2_-transformed median transcripts per mapped million reads (TPMs) of *F. hepatica plk* genes in *in vitro and in vivo* 3-week juveniles [46]. (C) Conservation of functional motifs in FhPLK1 but not FhPLK2/4 when compared to PLK1 sequences from *Echinococcus multilocularis*, *Schistosoma mansoni*, *Drosophila melanogaster*, *Homo sapiens* and *Mus musculus*; * = conserved residue, : = highly conserved amino acid properties, . = weakly conserved amino acid properties (our transcriptomic reads were used to confirm sequence).

### RNAi-mediated gene silencing of *fhplk1* disrupts juvenile growth

To test if *fhplk1* is integral to maintenance of the *F. hepatica* neoblast-like cell population we employed RNAi to probe its function *in vitro*. Parasites were treated with *fhplk1*-specific dsRNA and monitored for signs of abnormal growth. Transcript knockdown of *fhplk1* was measured after four weeks and found to be reduced by ∼90% relative to control treatment (Fig 3A; mean transcript expression ±SEM: control dsRNA = 127.67±9.49%, *fhplk1* dsRNA = 9.33±2.73%). We also found that transcript expression of another proliferative cell marker (histone 2b (*fhh2b*)) was reduced by 81.7±9%. Though not significantly different, it suggested that *fhh2b* expressing cells were depleted in *fhplk1-*RNAi worms; expression of a muscle cell marker (myosin light-chain (*fhmlc*)) was reduced by 21.3±30% relative to untreated worms, although again this was not significant (Fig 3A). H2B is highly expressed throughout the S-phase of the cell cycle when it participates in chromatin organisation [75]. Histone depletion reduces cell-cycle progression [76,77] and RNAi-mediated silencing in free-living and parasitic flatworms impairs proliferation, leading to a failure in tissue regeneration/maintenance [12,16,21,78]. Here, we observed that *fhplk1-*RNAi affected fluke development *in vitro*, with *fhplk1*-RNAi juveniles significantly smaller than control worms three weeks after the first RNAi trigger (Fig 3B; mean juvenile area ±SEM at three-weeks: untreated = 97350±2559 µm^2^, control-dsRNA = 94214±4266 µm^2^, *fhplk1* dsRNA = 73964±2559 µm^2^). This phenotype was further exaggerated after a fourth week (Fig 3B; mean juvenile area ±SEM at four-weeks: untreated = 118368±4732 µm^2^, control-dsRNA = 117781±5130 µm^2^, *fhplk1* dsRNA = 75942±2495 µm^2^). We also found that *fhplk1* knockdown impaired cell proliferation in four-week-old juveniles, thereby disrupting the neoblast-like stem cell population (Fig 3C & D; mean # EdU^+^ nuclei ±SEM: control-dsRNA = 477.5±47; *fhplk1*-dsRNA = 9.8±5.4). RNAi of *plk1* in mammalian cancer cell lines *in vitro* also reduced proliferation [79], and reduced tumour size *in vivo* in mice with non-small cell lung carcinoma [80], supporting a conserved role for *plk1* in proliferative cells. Interestingly, following four weeks of *plk1* silencing, juveniles cultured for a week in the absence of a RNAi-trigger showed no signs of recovery and did not resume proliferation (S2 Fig); this observation is consistent with the hypothesis that prolonged *fhplk1*-RNAi results in the ablation of *F. hepatica* neoblast-like stem cells. Together, these data support a conserved function for PLK1 in *F. hepatica* that ensures cell-cycle progression. As PLK1s are functionally conserved in *S. mansoni* and *E. multilocularis* [40,43], targeting PLK1 activity could be an effective pan-phylum control strategy.

**Fig 3.**
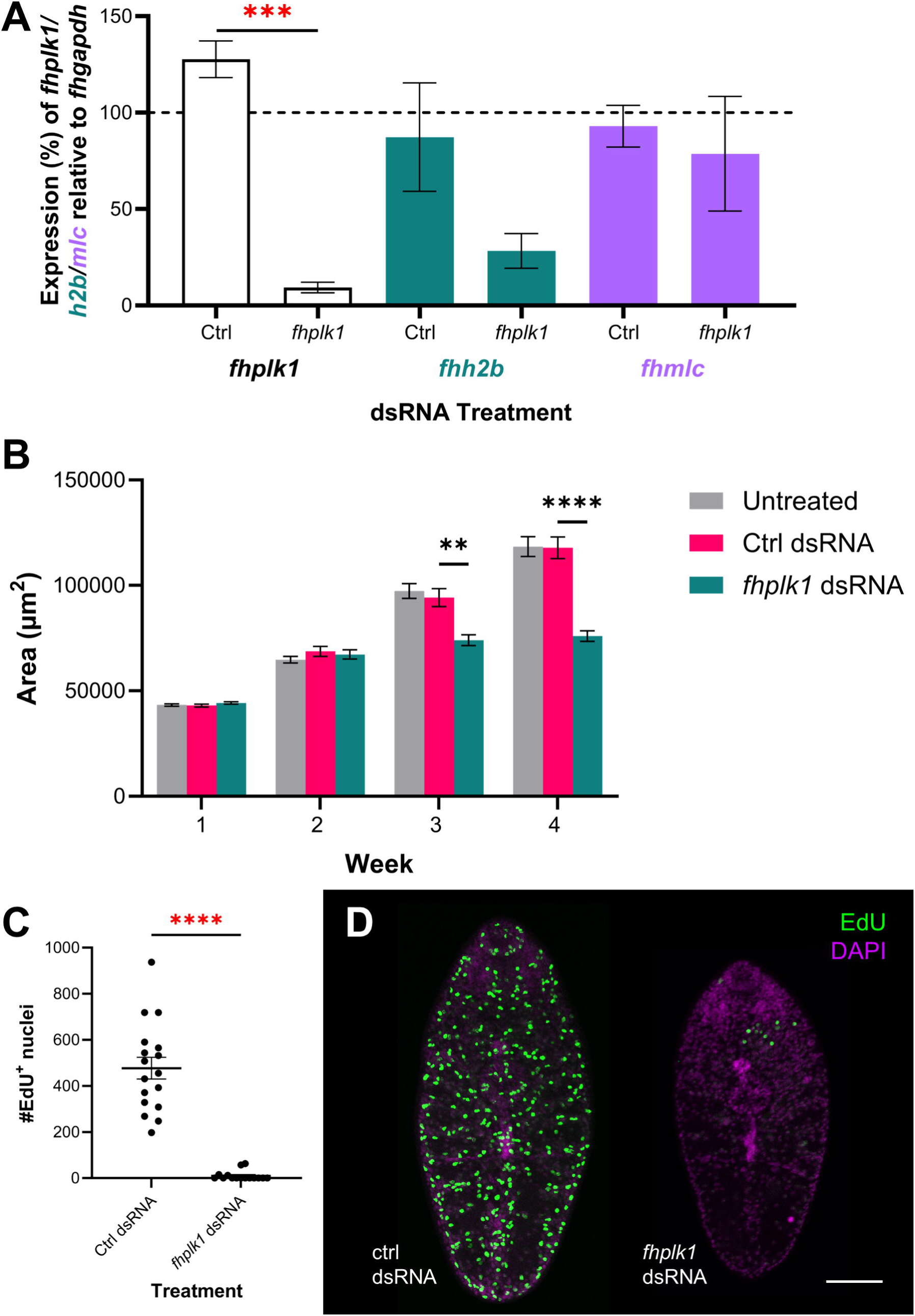
Knockdown of *fhplk1* disrupts growth and cell proliferation in juvenile *Fasciola hepatica in vitro*. (A) Mean expression ±SEM of *fhplk1* (white), *fhh2b* (green) and *fhmlc* (purple) relative to *fhgapdh* in juvenile *F. hepatica* following four weeks of repeated exposures to *fhplk1* dsRNA *in vitro* shows significant knockdown of *fhplk1* and reduction in expression of neoblast-like cell associated gene *fhh2b* relative to control dsRNA treated worms (unpaired t tests). (B) Mean area ±SEM of juvenile *F. hepatica* significantly diminished over three/four weeks of *fhplk1* dsRNA exposures (Kruskal-Wallis and Dunn’s posthoc tests). (C) Mean # EdU^+^ nuclei ±SEM significantly decreased after four weeks of *fhplk1* dsRNA treatments in juvenile *F. hepatica* (Mann-Whitney U test). (D) Confocal images of EdU staining (green) in four-week-old *F. hepatica* treated with *fhplk1* dsRNA *in vitro* shows reduction in # EdU^+^ nuclei compared to control dsRNA treated worms; scale bar = 100 µm, DAPI counterstain (magenta). **, p<0.01; ***, p<0.001; ****, p<0.0001.

### RNAi of *fhplk1* downregulates cell cycle effectors and ribosome biogenesis

We previously used irradiation to identify transcripts associated with *F. hepatica* neoblast-like cells [21]. However, while irradiation is effective at ablating proliferating cells, the potential for damage to differentiated cells limits its utility. Transcriptomic analysis following *fhplk1* silencing would help inform *fhplk1*’s role in juvenile fluke and support identification of other genes associated with growth/development. Therefore, we carried out target-specific RNAi for three weeks before worm extraction and transcriptome sequencing (Fig 4A), confirming that *fhplk1*-RNAi reduces growth and ablates neoblast-like stem cells (S3A & B Fig). RNA-Seq was then used to identify differentially expressed genes (DESeq2, padj < 0.001; S4 Table), with 946 downregulated genes and 1244 upregulated genes (Fig 4B). TOPGO analysis showed that genes associated with protein translation, ribosome assembly and DNA replication were overrepresented among downregulated genes (Fig 4C). The downregulation of genes associated with protein synthesis (‘translation’ and ‘protein folding’; Fig 4C) was interesting as studies in mouse cells showed that protein synthesis rates in early differentiated mouse cells are two-fold of that in embryonic stem cells [81], a result also observed in other mammalian cell cultures (reviewed in [82]). As *fhplk1*-RNAi depletes neoblast-like cells (i.e. stem cells) the proportion of differentiated cells might be expected to increase relative to control worms, although the production of more undifferentiated cells in the control groups could explain the higher levels of protein synthesis. Further, other GO terms (‘ribosomal small subunit assembly’, ‘structural component of ribosome’; Fig 4C & S3C Fig) suggested that the downregulation of translation may be related to reductions in ribosomal activity. KEGG pathway analysis showed the downregulation of ribosomal subunits/biogenesis following *fhplk1*-RNAi (S3D Fig) which matches observations in other platforms where stem cells have lower translation rates than differentiated cells, except in the case of ribosome biogenesis, which is upregulated in proliferating cells [83]. The downregulation of ribosome biogenesis-associated genes was also observed 14 days after irradiation in *S. mansoni* [13], demonstrating that this occurs in related species. It is also possible that the differential expression of ribosome-associated genes/pathways is indicative of general stress, as downregulation of these genes/pathways has previously been noted in nematodes following drug treatment (with non-stem cell targeting drugs) and local environment change [84,85].

**Fig 4.**
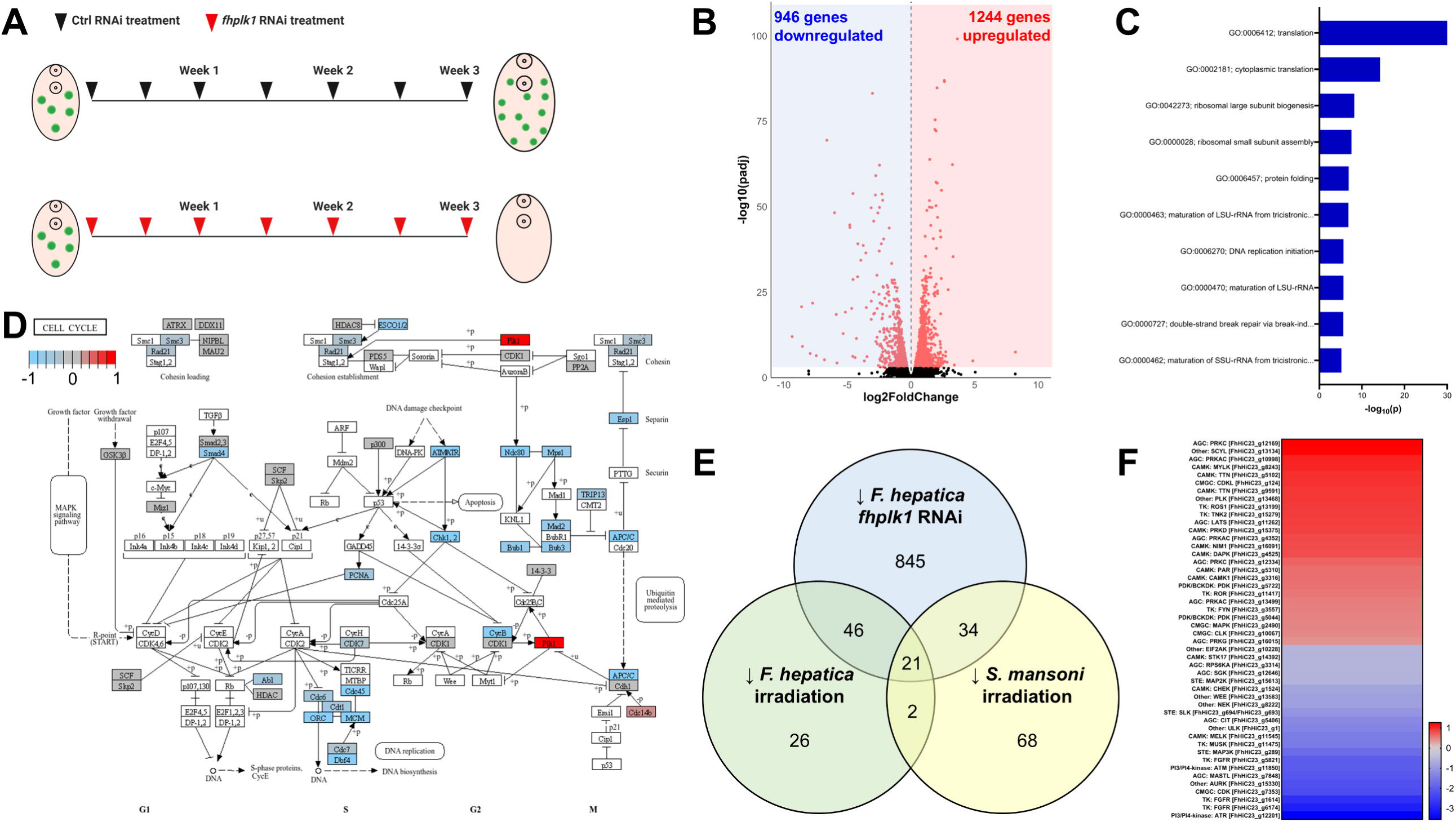
*fhplk1* knockdown downregulates cell cycle and translation-associated transcripts in juvenile *Fasciola hepatica in vitro*. (A) Experimental timeline showing dsRNA treatments (24 h dsRNA exposure each time) carried out on juvenile *F. hepatica* across three weeks to generate transcriptomes with final treatment from day 21-22. Volcano plot showing differentially expressed genes (red dots; DESeq2, padj < 0.001) after *fhplk1* RNAi in juvenile *F. hepatica* where blue shading indicates downregulated genes and red shading indicates upregulated genes. (C) Top 10 overrepresented biological process GO terms among downregulated genes following *fhplk1* RNAi shows genes involved in translation and nucleotide interactions. (D) Many components of KEGG cell cycle pathway downregulated following *fhplk1* RNAi (red = upregulated; grey = no change; blue = downregulated; white = unassigned KEGG ID). (E) Venn diagram showing overlap between genes downregulated following *fhplk1* RNAi in juvenile *F. hepatica*, irradiation in juvenile *F. hepatica* (green) or irradiation in adult *S. mansoni* (yellow). (F) Log_2_foldchange of differentially expressed kinases following *fhplk1* RNAi shows downregulation of proliferation-associated kinases such as MAPKs and FGFRs (colours correspond to Log_2_foldchange; red = upregulated; blue = downregulated).

Transcripts associated with DNA/RNA interactions and the cell cycle were also downregulated following *fhplk1* knockdown (e.g. DNA-replication initiation; Fig 4C & S3 Fig). Indeed, 4/5 KEGG pathways downregulated following irradiation of *F. hepatica* juveniles [21] were downregulated following *fhplk1* RNAi (S3D Fig). The absence of actively proliferating cells following *fhplk1*-RNAi, and concurrent downregulation of cell cycle-associated transcripts adds to the evidence that proliferative cells in the somatic tissue of juvenile/immature fluke are analogous to the neoblast-like stem cells of other parasitic flatworms [12,13,16]. Numerous core cell cycle genes were downregulated (S3 Table), including *atm* and *atr* kinases, *p53*, cell division control proteins, *cyclinB*, members of the mini-chromosone complex (*mcm2-7*) and origin recognition complex subunits (*orc2 & 3*). Additional core cell cycle regulators not annotated by eggNOG, but identified through BLAST searches, were also downregulated, such as a p53 homologue (FhHiC23_g6525). p53 is a tumour suppressor which can arrest the cell cycle and cause apoptosis when activated [86]. Mutations in p53 are linked to cancerous growths, with compounds developed to reverse these adverse mutation effects currently in clinical trials [87]. Furthermore, a *S. mansoni* p53 homologue has been linked to their neoblast-like cells [18]. While many of the cell cycle effectors discussed likely have conserved structure/function in both parasites and their hosts, inhibitors that indirectly target them [88] may be worth exploring for their anthelmintic potential. One surprising observation in the cell cycle schematic was ‘upregulation’ of *fhplk1* following *fhplk1* RNAi (DESeq2, *fhplk1* Log_2_Fold Change = ↑ 3.73). Our earlier trials had confirmed *fhplk1* knockdown following dsRNA treatment via qPCR (Fig 3A) so this upregulation was unexpected. However, examination of the read files via the Integrative Genomics Viewer (https://igv.org/) revealed that *fhplk1* dsRNA was likely sequenced as reads were observed only in the 192 bp dsRNA amplicon region of *fhplk1* dsRNA samples (S3E & F Fig). Control dsRNA samples on the other hand had reads across the length of the gene, with exons clearly visible (S3E Fig). The data indicate that this sequencing anomaly is related to our RNAi methodology as we ensure knockdown by repeatedly soaking worms in 100 ng/µL dsRNA) [89], including the 24 h period immediately preceding worm extraction.

We aimed to identify pan-trematode transcripts clearly linked to parasitic flatworm neoblast-like cells by cross-referencing the transcriptomic data generated here against the transcriptomes of irradiated *S. mansoni* and *F. hepatica* [12,21]. We found 21 genes downregulated in all three transcriptomic datasets following neoblast-like cell ablation (Fig 4E; note that the relatively small number of downregulated genes found in both irradiated datasets is likely due to the different life stages investigated as discussed in [21]). Many of the genes downregulated in all three datasets are known for their links to proliferation (S4 Table), including the cell cycle regulators *p53*, *cyclinB* and the *mcm* complex members. A further 34 genes were downregulated following *fhplk1*-RNAi in *F. hepatica* and irradiation in *S. mansoni*, including *hepatic leukaemia factor*, *mastL* (*greatwall kinase*) and *wdhd1* (*WD repeat and HMG-box DNA-binding protein/and-1*) which again have been associated with cell cycle regulation [90], DNA replication [91] and stem cell population maintenance [92]. Finally, 46 genes were downregulated in *F. hepatica* following either *fhplk1*-RNAi or irradiation with eight transcription factors/regulators included among them as well as an *fgfr*, implying conserved roles related to *F. hepatica* neoblast-like cells. While there were these shared downregulated genes between these two *F. hepatica* datasets there were 879 genes only downregulated following *fhplk1*-RNAi (Fig 4E). We hypothesise that this difference is caused by the loss of neoblast-like cells over an extended 3-week period of time (unlike the irradiated dataset in which juveniles were sequenced 48 hours after X-ray exposure), with this having knock-on effects across tissues in the worm.

As several of the genes downregulated in both datasets were kinases we explored the expression all *F. hepatica* kinases following *fhplk1* RNAi; 48 kinases were differentially expressed (22 downregulated and 26 upregulated; Fig4 F). Among downregulated kinases were three *fgfrs* that were also upregulated in developing life stages (FhHiC23_g6174, FhHiC23_g1614 and FhHiC23_g5821), an aurora kinase and members of the MAPK signalling cascade which regulates cell cycle progression [93]. Again, these kinases are integral to activate proliferation in other species and may be appealing targets for control. AGC kinases and CAMKs were more represented among upregulated kinases, though their functions are more diverse and less clearly linked to proliferation.

One final point to note is that of 46 genes downregulated after irradiation or *fhplk1*-RNAi in *F. hepatica*, ten had no clear BLAST hit in model species, though they all have *S. mansoni* homologues and most have *F. gigantica* homologues. Furthermore, of the 50 most downregulated transcripts (ranked by fold-change) only 8 had clear BLAST hits in model species (e-value <0.001, S3 Table), though most were found to have orthologues in at least one other parasitic flatworm. While we know little of the function of these genes/proteins, their downregulation links them to putative roles in *F. hepatica* growth/development associated with neoblast-like stem cells, especially where also downregulated following irradiation. These unknown genes could be attractive drug targets, given their lack of homology to genes in host species.

### Inter-cell signalling upregulated following *fhplk1* RNAi

Over 1200 juvenile fluke genes were upregulated following three weeks of *fhplk1*-RNAi (Fig 4B; S3 Table). TOPGO analysis of these genes showed that GO terms associated with inter-cell signalling systems such as ion transport, cell-adhesion and synapses were overrepresented (Fig 5A; S4A Fig) among the upregulated genes. Furthermore, cellular component GO terms suggested many predicted proteins are associated with the cell membrane and extracellular space/synapse (S4A Fig), supporting evidence for the upregulation of inter-cell signalling. KEGG pathway analysis further reinforced this, highlighting upregulation of multiple neuronal-ligand receptors, including many GPCRs (Fig 5B). Many signalling pathways were also upregulated, such as those linked to the gap junction as well as GABAergic, serotonergic and cholinergic synapses (S4B Fig). Gap junctions are critical to cell-cell connectivities that help coordinate cellular functions in multicellular organisms, though invertebrates lack the associated connexin proteins and instead possess innexins [94]. While the gap junction itself is unmarked in the KEGG pathway of S4C Fig as the pathway is based on that of humans, we found five predicted innexins to be significantly upregulated following *fhplk1*-RNAi (FhHiC23_g648, FhHiC23_g6222, FhHiC23_g9940, FhHiC23_g15433 and FhHiC23_g15004; S3 Table). Innexins have been localised in various planarian tissues, including the nervous system [95], and their inhibition can lead to developmental abnormalities through polarity disruption during regeneration [95,96], or ablation of neoblast-cell populations [97], suggesting key roles in developmental regulation.

**Fig 5.**
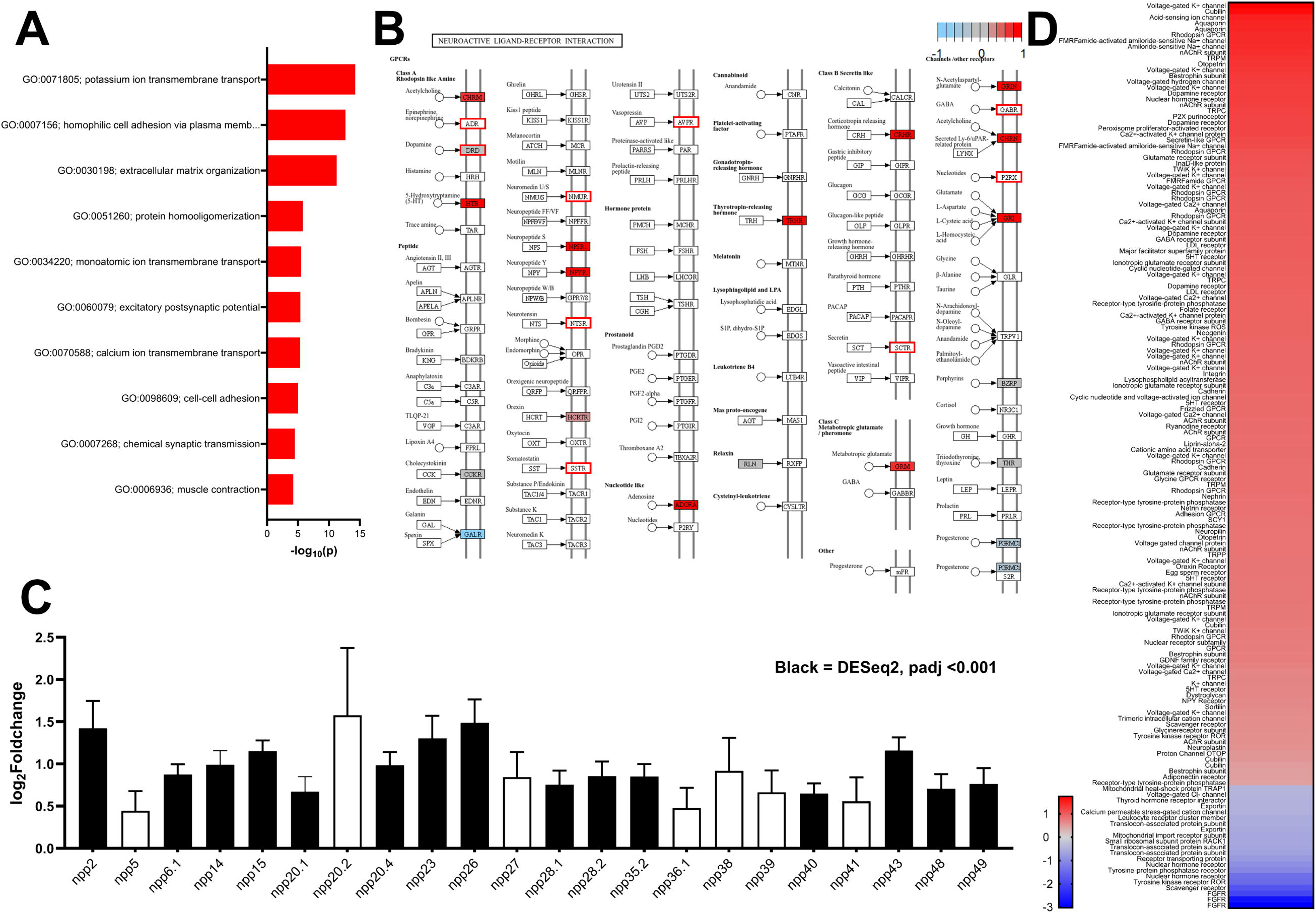
RNAi of *fhplk1* in *Fasciola hepatica* results in upregulation of inter-cell signalling systems. (A) Biological process GO terms (top 10) associated with ion transport and cell signalling are overrepresented among upregulated genes following *fhplk1* RNAi. (B) KEGG pathway showing upregulation of several neuroactive-ligand receptors following *fhplk1* RNAi (red = upregulated; grey = no change; blue = downregulated; white = unassigned KEGG ID; red box = unassigned KEGG ID but identified as upregulated from BLAST searches). (C) Log_2_foldchange ±SEM of putative *F. hepatica* neuropeptides following *fhplk1* RNAi shows all 22 upregulated, with 15 differentially expressed (black bars). (D) Log_2_foldchange of differentially expressed putative receptors and channel subunits (identified via BLAST searches) following *fhplk1* RNAi shows upregulation of 134/155 (colours correspond to Log_2_foldchange; red = upregulated; blue = downregulated).

Within the upregulated classical transmitter pathways there was upregulation of ligand synthesis enyzmes, receptors and downstream signalling molecules such as G proteins (GABAergic example shown in S4D Fig). Indeed, of the known *F. hepatica* classical transmitter pathways [98], only the histamine pathway had no evidence (based on TOPGO, KEGG and BLAST analysis) of upregulation, a poorly understood pathway in parasitic flatworms [99]. Curiously, some upregulated classical transmitter pathways have opposing effects on *F. hepatica* motility (e.g. acetylcholine is believed to be inhibitory [100], while serotonin is excitatory [101]), making their concurrent upregulation interesting and potentially linked to increased tonic regulation. Serotonin has been implicated in proliferation in other species [102–105] and has been shown to directly stimulate growth/proliferation in *S. mansoni* and *E. multilocularis* [106,107]. While the roles of acetylcholine and dopamine in flatworm growth are unknown, they have been linked to proliferation in mammals [108,109]. Furthermore, dopamine deficiency in *C. elegans* and *D. melanogaster* constrains development [110,111] and a dopamine antagonist reduced *S. mansoni* miracidial transformation [112]. *C. elegans* lacking choline acetyltransferase (ChAT) also display slower growth [113].

Additionally, 16/22 putative neuropeptide encoding genes identified in our dataset (S5 Table) were significantly upregulated in *plk1*-RNAi worms (Fig 5C), as were their processing enzymes; *prohormone convertase 2* (*pc2*; FhHiC23_g9059), *carboxypeptidase E* (*cpe*; FhHiC23_g10997), *peptidylglycine alpha-hydroxylating monooxygenase* (*phm*; FhHiC23_g3039) and *peptidyl-alpha-hydroxyglycine alpha-amidating lyase* (*pal*; FhHiC23_g9107), along with several predicted neuropeptide receptors (Fig 5B). Neuropeptides have been shown to drive mammalian cell proliferation [114], as well as germline proliferation and regeneration in planarians [115–118]. Furthermore, two neuropeptides and their GPCRs are required for normal maturation of female reproductive structures in *S. mansoni* [119]. If classical transmitters and neuropeptides have conserved roles in managing growth/proliferation in *F. hepatica,* then the upregulation observed could indicate that *fhplk1*-RNAi juveniles lacking neoblast-like cells are attempting to boost cell proliferation via enhancement of selected inter-cell signalling systems.

Previously, we highlighted that signalling systems were upregulated in slower growing *F. hepatica* with decreased cell proliferation [46]. Here we saw the loss of neoblast-like stem cells, concomitant with nervous system upregulation, indicating potential interplay between proliferating neoblast-like stem cells and neuronal signalling systems. When we compared upregulated genes in our neoblast-like stem cell ablated juveniles with downregulated genes in neoblast-like stem cell-enriched *in vivo* juveniles [46], we identified 244 shared genes, including 12 neuropeptide encoding-genes (S6 Table). Further, TOPGO analysis of these genes showed that neurotransmitter and Wnt signalling were enriched (S4E Fig). Enrichment of the Wnt pathway is notable as Wnts support proliferation and polarity orientation in regenerating planarians [120], with similar roles proposed in parasitic flatworms [121–123]. RNAi of *wnt* genes in regenerating planarians can result in aberrant neurogenesis [124–127] and their silencing was recently shown to inhibit *F. hepatica* neoblast-like cell proliferation and growth/development *in vitro* [128].

To help us understand the scale of signalling upregulation we examined all BLAST results for the terms ‘receptor’ or ‘channel’ to identify putative intercellular signalling-related genes. Of the 155 putative receptors/channels identified through BLAST searches, 134 (86%) were significantly upregulated following *plk1*-RNAi (Fig 5D). Those that were downregulated were mainly kinases or nucleus-associated transporters that could be associated with worm neoblast-like stem cells. S4G Fig shows that 22% of the upregulated receptors/channels were predicted rhodopsin peptide GPCRs with voltage-gated potassium channel subunits constituting another large component. Among these genes were two putative FMRFamide-activated amiloride-sensitive Na+ channel subunits [129] that may be of interest as putative drug targets due to their unusual pharmacology and proposed absence in mammals. These data illustrate the extent of upregulation seen within signalling systems following sustained *fhplk1*-RNAi.

We recognise that the reduced size of *plk1*-dsRNA worms (in comparison to control dsRNA worms) may increase the proportionality of neuronal tissue, thereby increasing the balance of neuronal-associated transcripts. However, tissue proportionality does not appear to be a key factor in the observed upregulation for a variety of reasons. Firstly, while *fhplk1*-RNAi worms were indeed smaller (S3A Fig), the difference in size was modest compared to that seen in the gene-function RNAi trials (Fig 3B). Furthermore, while we found DAPI staining (calculated as a percentage of worm area) decreased following *fhplk1*-RNAi (S4F Fig), the decrease was ∼20% which may be directly attributable to the absence of neoblast-like stem cells (neoblasts account for around 20-35% of all cells in free-living flatworms [130]). Second, another study observed the downregulation of signalling systems in faster growing, stem cell enriched *F. hepatica* [46], with much greater size differences (∼15x) than those observed in our study. Despite this, genes associated with signalling had comparable fold changes in expression in most instances (S7 Table), again suggesting size/tissue proportionality is not a factor in nervous system upregulation. Thirdly, we did not observe consistent upregulation of transcripts associated with other tissue types, such as the tegument (*tsp2*) or the gut (18 cathepsins upregulated and 14 downregulated) which indicates that the nervous system transcript upregulation is unusual. Finally, we found this pattern of increased signalling in adult male *S. mansoni* datasets in which worms were subjected to transcriptomics two weeks after irradiation, or three weeks after the onset of *smfgfrA* or *smh2b* RNAi [13]. TOPGO analysis of these datasets showed again that there was an increase in transcripts associated with signalling systems, ion transport and cell communication (S8 Table). Given that these worms were adults of comparable size and that the anterior portion, including cerebral ganglia, had been removed (to exclude testes from analysis) the upregulation of signalling systems following neoblast-like stem cell ablation in diverse circumstances supports the validity of these transcriptional changes in neuronal signalling systems and warrants further investigation.

These data challenge the traditional orthodoxy of using short-term motility-based assays to assess the drug target candidature of various helminth signalling pathways. If the signalling systems highlighted in this study do play significant roles in cell proliferation, then potential long-term impacts of drugs on worm growth/development could be missed with restricted phenotype monitoring windows. Glutamate-gated chloride channels and nicotinic acetylcholine receptors are already targets of current anthelmintics, while GPCRs are thought of as highly ‘druggable’ targets [131]. Repurposing drugs to disrupt neoblast-like stem cell biology could expose novel control strategies.

### No correlation between DE miRNAs and predicted DE mRNA targets

We carried out miRNA sequencing on the same RNA samples and used miRDeep2 to map reads to the 150 predicted miRNAs [132]. We found 68 of these miRNAs were expressed in at least two of the six samples, with 13 downregulated and 8 upregulated (S5A Fig; DESeq2, padj <0.001; S9 Table). Target prediction software was then used to identify putative target transcripts of these differentially expressed miRNAs (S9 Table). TOPGO analysis did not identify any enriched GO terms among potential target transcripts. We then plotted GO term frequency of predicted target transcripts against fold change (S5B Fig) but found little, to no overlap with GO terms of differentially expressed mRNAs. Finally, only 36 targets predicted to be downregulated were found to be so, while only 38 of those predicted to be upregulated were so. No clear links between *fhplk1*-RNAi silencing in juvenile *F. hepatica* and endogenous miRNA activity were uncovered.

### Pharmacological inhibition of FhPLK1 phenocopies *fhplk1*-RNAi

We next assessed the drug-target candidature of FhPLK1 through use of the commercially available inhibitor BI 2536. Overexpression of PLK1 in various cancers has led to the development of inhibitors, including BI 2536, a highly selective PLK1 inhibitor which can blunt cell proliferation in various cancer types, both *in vitro* and *in vivo* [133,134]. As BI 2536 is an ATP-competitive inhibitor [133], high levels of sequence similarity across human PLK1-3 kinases compromises its selectivity [133,134]. As discussed previously we identified two additional PLK genes in *F. hepatica*, one similar to human PLK2 (FhPLK2), with the other similar to both human PLK4 (FhPLK4) and SAK (Serum-inducible kinase Akin Kinase) in *S. mansoni* and *E. multilocularis* [43,135]. PLK4 divergence in humans renders it less sensitive to BI 2536 [133], a trait also observed for *S. mansoni* SmSAK [135]. Crystallisation experiments identified Leu132 of HsPLK1 as a key residue for facilitating drug-target interaction and selectivity. This leucine was conserved in FhPLK1 (Leu104), but not in FhPLK2 or FhPLK4 (S6A Fig), suggesting that the activity of BI 2536 would be specific to FhPLK1 in *F. hepatica*.

Initial experiments involved exposing NEJs to a broad range of BI 2536 concentrations (0.001 µM – 10 µM) for 18 hours in RPMI, with motility measured as an indicator of viability as motility of *F. hepatica* had previously been disrupted following BI 2536 exposure [45]. However, in our hands, none of the BI 2536-treated groups exhibited reduced motility (S6B Fig). We monitored rhythmic movement (length of individual parasites plotted across frames) for subtle motility changes (S6C Fig), finding that even those treated with 10 µM BI 2536 retained the classical, ‘accordion’-style movement associated with *F. hepatica* NEJs (rhythmical lengthening and shortening in a probing fashion) [136]. This was in stark contrast to treatment with 1 µM TCBZ which paralysed juveniles (S6C Fig). Our BI 2536 results are not directly comparable to those of Morawietz et al. [45] as they treated older worms (*ex vivo* immature and adults) with higher concentrations (20 µM – 100 µM).

As neoblast-like cell inhibition is associated with growth defects [20,21], worms were treated with BI 2536 for 18 h and then cultured for one week in the absence of drug. There were no significant differences between the sizes of juveniles from any treated group and the DMSO-treated controls (Fig 6A). However, incubation of a subset of juveniles with EdU showed significantly reduced proliferation in those treated with 10 µM BI 2536 (Fig 6B; # EdU^+^ nuclei ±SEM: DMSO = 465.1±109.5, 10 µM BI 2536 = 55.1±25.9). A small reduction in EdU^+^ nuclei was also seen in juveniles treated with 1 µM BI 2536, though this difference was not significant. These data suggest that culture following drug exposure afforded the opportunity for fluke to recover. To test this, parasites were cultured for one week before treatment with BI 2536 and subsequent EdU exposure. These worms displayed concentration-dependent reduced cell proliferation across all groups, with significance observed at both 1 µM and 10 µM BI 2536 (Fig 6C; # EdU^+^ nuclei ±SEM: DMSO = 173.8±11.5, 1 µM BI 2536 = 12.7±4.7, 10 µM BI 2536 = 0±0). The observed phenotypic responses are consistent with BI 2536 inhibiting PLK1 activity in *F. hepatica*, matching observations in other parasitic flatworms [40,43].

**Fig 6.**
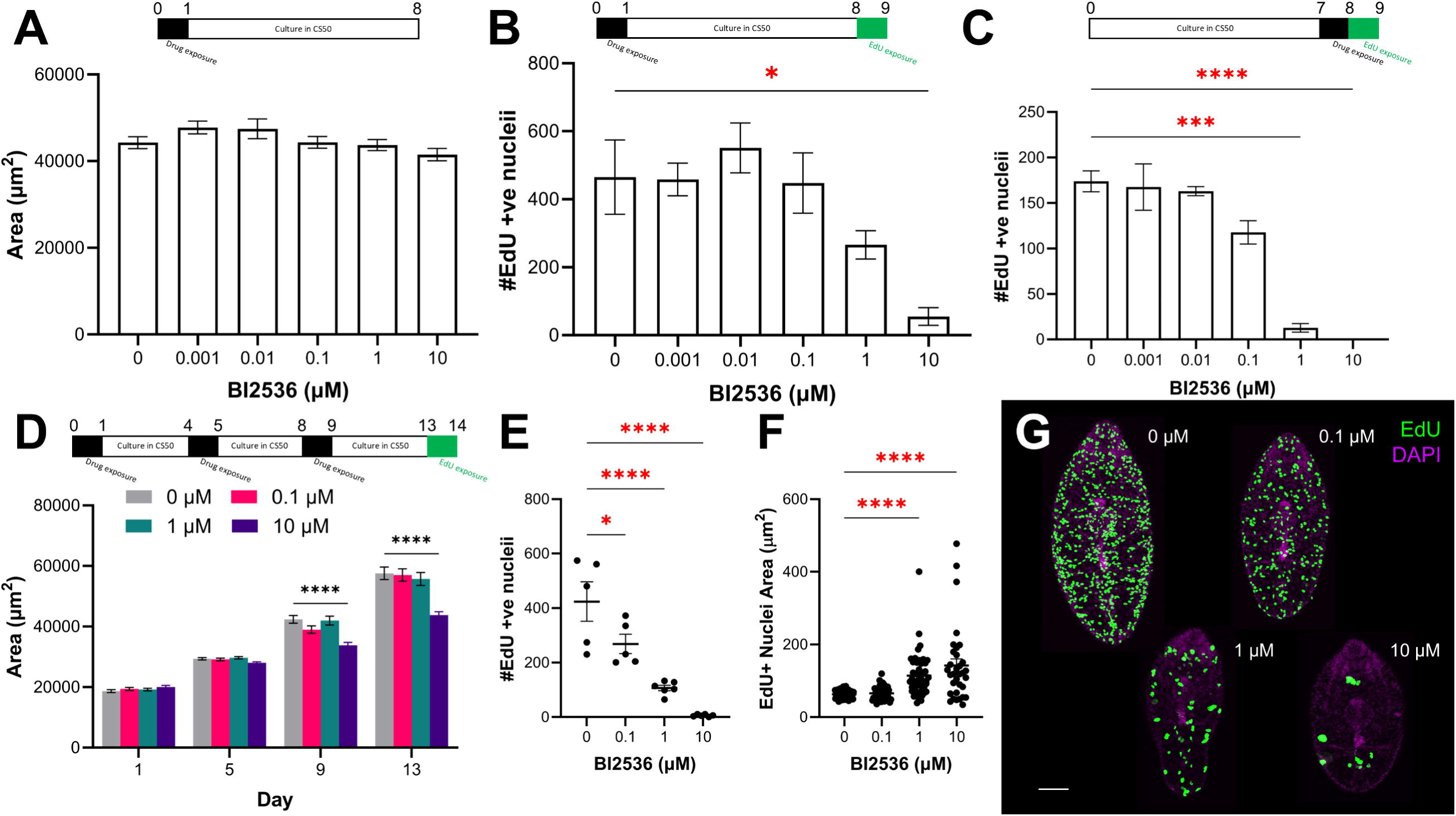
Repeated exposure of juvenile *Fasciola hepatica* to the PLK1 inhibitor BI 2536 *in vitro* reduces growth and proliferation. (A) Mean worm area ±SEM of 7-day-old juvenile *F. hepatica* after exposure to BI 2536 on day 0, followed by 7 days in culture, shows no effect on growth. (B) Mean # Edu^+^ nuclei ±SEM in 9-day-old *F. hepatica* after exposure to BI 2536 on day 0, followed by 8 days in culture, shows 10 µM significantly reduced the number of proliferating cells. (C) Mean # Edu^+^ nuclei ±SEM in 9-day-old *F. hepatica* after exposure to BI 2536 on day 7, shows both 1 µM and 10 µM significantly reduced the number of proliferating cells. (D) Mean area (µm^2^) ±SEM of *F. hepatica* across 13 days following BI 2536 exposures on days 0, 4 and 8 shows 10 µM significantly reduced growth rate. (E) Mean # Edu^+^ nuclei ±SEM in 14-day-old *F. hepatica* after repeated BI 2536 exposures on days 0, 4 and 8 (timeline from (E) applies to this graph) shows all tested concentrations significantly reduced the number of proliferating cells. (F) Mean size of Edu^+^ nuclei ±SEM in 14-day-old *F. hepatica* after repeated BI 2536 exposures (1 and 10 µM) on days 0, 4 and 8 (timeline from (E) applies to this graph) shows increased nuclei area. (G) Confocal images of EdU staining (green) in 14-day-old *F. hepatica* exposed to various concentrations of BI 2536 on days 0, 4 and 8 shows a dose-dependent reduction in # EdU^+^ nuclei and increase in EdU^+^ nuclei area compared to DMSO-treated control; scale bar = 100 µm, DAPI counterstain (magenta). *, p<0.05; ***, p<0.001; ****, p<0.0001.

As BI 2536 disrupted neoblast-like cell proliferation we investigated whether it could inhibit juvenile development. For these experiments, parasites were treated with BI 2536 multiple times (on days 0, 4 and 8) with measurements taken on days 1, 5, 9 and 13. After 9 days, worms treated with 10 µM were significantly smaller than control worms (Fig 6D; day 9 mean juvenile area ±SEM: DMSO = 42368±1286 µm^2^, 10 µM BI 2536 = 33861±907 µm^2^). This phenotype was still evident following a drug-free period post treatment (Fig 6D; day 13 mean juvenile area ±SEM: DMSO = 57568±2035 µm^2^, 10 µM BI 2536 = 43799±1122 µm^2^). EdU exposure revealed significant, concentration-dependent reductions in cell proliferation across all groups (Fig 6E & G; # EdU^+^ nuclei ±SEM: DMSO = 423.8±72.3, 0.1 µM BI 2536 = 268.6±35.5, 1 µM BI 2536 = 107±10, 10 µM BI 2536 = 6±1.8). These results largely phenocopy those seen following *fhplk1*-RNAi (Fig 3), supporting FhPLK1’s conserved regulatory role in the proliferation of *F. hepatica* neoblast-like stem cells. We noticed that EdU^+^ nuclei appeared larger in worms treated with higher concentrations of the drug. Therefore, we measured 10 randomly selected EdU^+^ nuclei from five juveniles in each treatment group. This revealed that the EdU^+^ nuclei were significantly larger in worms treated with 1 µM and 10 µM BI 2536 (Fig 6F & G; EdU^+^ nuclei area: DMSO = 63.2±1.6 µm^2^, 1 µM BI 2536 = 114.3±8.2 µm^2^, 10 µM BI 2536 = 142.1±18.1 µm^2^). These larger nuclei could be indicative of neoblast-like cells that had passed through S phase before being delayed/arrested at either the G2/M transition or during mitosis. BI 2536 has previously been shown to increase the DNA content in cancer cells, which was attributed to an inability to undergo cytokinesis [134]. We did not observe larger EdU^+^ nuclei throughout our RNAi experiments. This could be due to the short-term effects of inhibitor exposure which could be reversed following its removal in contrast to the long-term systemic changes to the biology of the worms throughout *fhplk1* RNAi [137].

While BI 2536 has previously been shown to affect *F. hepatica* movement *in vitro* [45], our *plk1*-RNAi and pharmacological inhibition experiments highlight how disrupting FhPLK1 activity can undermine neoblast-like cell mediated growth and development. This could weaken *F. hepatica* juveniles within the host environment, aiding parasite clearance by the immune system. BI 2536 has proven anthelmintic activity against both *S. mansoni* and *E. multilocularis*, disrupting gamete production and preventing metacestode formation from germinative cells, respectively [40,43]. Taken together, these data suggest that developing/repurposing drugs to target PLKs could be a promising anthelmintic strategy to control a range of parasitic flatworms, including different life stages.

## Conclusion

This study encourages exploration of parasitic flatworm neoblast-like cells as targets for novel anthelmintics. Our annotation of the *F. hepatica* kinome showed that many kinases linked to proliferation are enriched in the highly pathogenic immature life stages. Functional genomics and inhibitor trials validated this approach as disruption of *fhplk1*, a core cell cycle-related kinase, diminished proliferation in the neoblast-like stem cell population and undermined worm growth/development. This approach could be expanded to other kinase targets such as the additional putative PLKs or the FGFRs. Indeed, the *S. mansoni* SmSAK gene has conserved roles in cell cycle progression [54], and knockdown reduced schistosomula viability *in vitro* [33]. Moreover, the ability of a kinase inhibitor to disrupt *F. hepatica* growth/development adds credence to the idea of targeting parasitic flatworm kinases [138], perhaps even in combination with current flukicides, to improve efficacy in cases of reduced drug susceptibility/resistance.

Our transcriptomic analyses highlighted possible links between *F. hepatica* neoblast-like cells and neuronal/signalling systems. Neurotransmitter signalling systems have long been touted, and exploited, as anthelmintic targets. However, if novel roles for these systems in parasitic flatworm growth/development are uncovered it opens new paths to target discovery and exploitation for drug development. A significant panel of genes were downregulated following *fhplk1*-RNAi, suggesting they have some role in mitotic regulation/stem cell biology in fluke. The work here has built new understanding of neoblast-like cell directed growth/development in *F. hepatica* which will enhance the opportunities for novel control target discovery.

## Methods

### Identification of putative *F. hepatica* protein kinases

The PK sequences of the nine *Homo sapiens* ePK families, along with the aPKs, were downloaded from Uniprot (https://ftp.uniprot.org/pub/databases/uniprot/current_release/knowledgebase/complete/docs/pkinfam.txt). HMMER (v3.4; http://hmmer.org/) was used to assemble query sequences for each ePK family and the aPKs. HMM searches were carried out using HMMER against the University of Liverpool and Washington University *F. hepatica* predicted proteomes [47,139] downloaded from Wormbase ParaSite (WBPS18, https://parasite.wormbase.org) [140]. We also looked for additional kinases in a set of novel transcripts. These were generated by aligning all publicly available *F. hepatica* transcriptomic reads [21,46,47] and in-house transcriptomic datasets against the University of Liverpool genome (WBPS19, https://parasite.wormbase.org/Fasciola_hepatica_prjeb58756/Info/Index/) using HISAT2 (v.2.1.0), before transcript assembly with Stringtie (v1.3.6) to create a master reference gtf file containing 14,684 ‘novel’ transcripts that did not overlap with genes in the reference assembly. All putative hits were screened via BLAST searches against the SwissProt database, and domains annotated using InterPro (https://www.ebi.ac.uk/interpro/about/interproscan/). Hits were retained only when the top BLAST result was a PK, and when a kinase domain was predicted to be present. Duplicates/putative gene copies were identified using Clustal Omega (https://www.ebi.ac.uk/Tools/msa/clustalo/) and manually inspected before removal. The remaining putative *F. hepatica* PKs were used to create two HMM query sequences (one ePK and one aPK) which were then used in further HMM searches against both *F. hepatica* predicted proteomes [47,139] to ensure no PKs were missed. Resulting hits were sorted through the same pipeline. BLAST results were used to classify which family/subfamily putative *F. hepatica* kinases belonged to.

Putative ePKs were clustered using CLANs analysis software (1,000,000 rounds of clustering; [141]). MEGA11 (https://megasoftware.net/, [142]) was used to align aPKs (ClustalW), identify the optimum model for phylogenetic analysis, and construct a Maximum Likelihood (ML) tree with 500 bootstraps. This ML tree was visualised using the Interactive Tree of Life (https://itol.embl.de/).

Published life-stage transcriptomic data from eggs, metacercariae, 1 h newly excysted juveniles (NEJs), 3 h NEJs, 24 h NEJs, three-week juveniles and adults [47] were used to obtain median TPM values for all predicted PKs. Z scores were generated, and heatmaps produced using Morpheus software (https://software.broadinstitute.org/morpheus).

### *F. hepatica in vitro* excystment and culture

The Italian strain of *F. hepatica* (Ridgeway Research, UK) was used for all experiments. Excystments were carried out as previously described [89,143], and a detailed protocol is available at https://dx.doi.org/10.17504/protocols.io.14egn212qg5d/v1. Briefly, metacercariae were physically ‘popped’ from their outer cyst walls and treated with 10% (v/v) sodium hypochlorite (#1056142500, Sigma-Aldrich) diluted in double-distilled water (ddH2O) for 2-3 minutes (exact time dependent on metacercariae batch). Popped and bleached metacercariae were then washed in ddH2O at least five times to remove excess sodium hypochlorite, prior to being stimulated to excyst as detailed by McVeigh et al. [143]. Parasites were cultured in 200 µL CS50 (50% (v/v) chicken serum (#16110082, Thermo Fisher Scientific) in RPMI (#11835105, Thermo Fisher Scientific) with antibiotic/antimycotic (#A5955, Sigma Aldrich)) in 96-well, round-bottomed plates (#83.3925, Sarstedt) in a humidified incubator at 37°C with 5% CO2 as previously described [20]. Media changes were performed three times per week. *F. hepatica* in culture media are referred to as juveniles.

### RNAi of fhplk1 in vitro

We generated cDNA from four-week-old juvenile *F. hepatica* by snap freezing 20 juveniles in liquid N2. Tissues were lysed in a Qiagen tissue lyser with a stainless-steel bead (50 oscillations/sec for 1 min) prior to mRNA extraction with the Dynabeads mRNA Direct Purification Kit (#61011, Thermo Fisher Scientific), DNase treatment (Turbo DNA-free Kit, # AM1907, Thermo Fisher Scientific) and reverse transcription to cDNA using the High-Capacity RNA-to-cDNA Kit (#4387406, Thermo Fisher Scientific). All cDNAs were diluted 1:1 in ddH2O prior to use. Double-stranded (ds)RNA templates specific to *fhplk1* (FhHiC23_g6132) were generated using 0.4 µM T7-labelled primers (5’-TAATACGACTCACTATAGGGT-3’) and the FastStart Taq DNA Polymerase, dNTPack (#4738357001, Millipore Sigma). Primers were designed using Primer3Plus [144], and amplicons were sequenced by Eurofins Genomics (https://eurofinsgenomics.eu/en/) to ensure target specificity. Primers used to generate *fhplk1*-specific dsRNA templates can be found in S10 Table. Negative control dsRNA targeting bacterial neomycin phosphotransferase [U55762] was generated as previously detailed [143]. Template sizes were checked on a 1-2% agarose gel prior to purification using the ChargeSwitch PCR Clean-Up Kit (#CS12000, Thermo Fisher Scientific) and dsRNA generation using the T7 RiboMAX Express RNA System (#P1700, Promega). All dsRNAs were resuspended in ddH2O with concentrations/purities checked on a DeNovix DS-11 FX spectrophotometer prior to storage as single-use aliquots at -20°C. RNAi treatments were performed by soaking worms in 50 µL of 100 ng/µL dsRNA diluted in RPMI for 24 h in 96-well round-bottomed plates under standard conditions detailed above. NEJs were immediately exposed to target-specific dsRNA with juveniles subsequently treated twice a week across four weeks to ensure knockdown [89]. Juveniles were cultured in CS50 between RNAi treatments as standard. All RNAi experiments were carried out in triplicate and included no dsRNA-treatment controls.

Juveniles were imaged each week by capturing darkfield videos/images on an Olympus SC50 camera attached to an Olympus SZX10 microscope. Video analyses were performed using the wrMTrck plugin (http://www.phage.dk/plugins/wrmtrck.html, [145]) for ImageJ calibrated to a 1 mm scale. The wrMTrck plugin was used as default except for modifications to minSize (10 pixels), maxSize (1000 pixels), maxVelocity (1000 pixels/frame), AreaChange (200% change in area between frames), Rawdata (2) and benddetect (0). Motility was measured by calculating changes in juvenile length between frames, which provided length change (µm)/minute. Movement of individual juveniles were normalised against mean movement of control juveniles. wrMTrck also provided area (µm^2^) measurements for each worm.

At the conclusion of RNAi trials juveniles were frozen in liquid N2 in batches of 20 worms/replicate before cDNA extraction as described above. Quantification of target transcript knockdown was carried out on a Qiagen RotorGene Q 5-plex HRM instrument under the following running conditions: 95°C for 10 minutes; 40 cycles @ 95°C for 10 sec, 55°C for 15 sec and 72°C for 20 sec. Melt-curve analyses were enabled to confirm product specificity and qPCR reactions were performed in triplicate using the SensiFast SBYR No-ROX kit (#BIO-98005, Bioline) with final primer concentrations of 200 nM. Glyceraldehyde phosphate dehydrogenase (*fhgapdh*) [AY005475] was used as a housekeeper gene. Alongside *fhplk1*, *fhh2b* [D915_007751] and *fhmlc* [FhHiC23_g308] were amplified as markers for proliferating cells and muscle cells respectively. The oligonucleotide primers used can be found in S10 Table. Pfaffl’s Augmented ΔΔCt method [146] was used to calculate relative gene expression. Transcript expression was plotted for both target and control-dsRNA-treated groups relative to no dsRNA control.

### Visualisation of proliferative cell activity

The thymidine analogue, 5-ethynyl-2-deoxyuridine (EdU; Thermo Fisher Scientific) was used to label nuclei of proliferating neoblast-like cells. A 10 mM stock of EdU was stored at -20°C in PBS. Stock solution was diluted in CS50 to a final working concentration of 500 µM and placed on juveniles for 24 h under standard culture conditions. Immediately following EdU exposure worms were flat-fixed under a coverslip in 4% paraformaldehyde (w/v) in PBS (PFA) for 15 minutes, prior to a further 4 h free-fixing in 4% PFA at room temperature while constantly rotating. Worms were permeabilised in 0.5% Triton X-100 in PBS for 30 min at room temperature. EdU detection was achieved using the Click-iT EdU Proliferation Kit for Imaging with the Alexa Fluor 488 dye (#C10337, Thermo Fisher Scientific). Worms were counterstained using 4′,6-diamidino-2-phenylindole (#D1306, DAPI) in PBS for 20 minutes at room temperature prior to mounting in Vectashield (#H-1000-10, Vector Laboratories). A detailed protocol is available at https://dx.doi.org/10.17504/protocols.io.eq2lyjnrrlx9/v1.

Samples were imaged on a Lecia TCS SP8 inverted microscope. Z-stacks consisting of 10-15 optical sections between the dorsal and ventral surfaces were captured for each worm and maximally projected for analyses. EdU^+^ nuclei counts were performed using the cell counter plugin for ImageJ (https://imagej.nih.gov/ij/plugins/cell-counter.html). Nuclei area (µm^2^) measurements were also performed in ImageJ, where images were calibrated to a 50 µm scale.

### RNA-seq and miRNA-seq of *fhplk1* RNAi *F. hepatica* juveniles

NEJs were excysted and treated with target-specific dsRNA as described above (∼50 worms/replicate split across two wells). Juveniles were treated with dsRNA in RPMI for the final time on day 21 before being snap frozen in liquid N2 the following day and stored at -80°C. Total RNA was extracted using Trizol reagent (#15596026, Thermo Fisher Scientific) before being sent to the Genomics Core Technology Unit at Queen’s University Belfast (https://www.qub.ac.uk/sites/core-technology-units/Genomics/) for RNA quantification/purity assessment on an AATI fragment analyser prior to library generation using the KAPA mRNA HyperPrep Kit (#KK8580, Roche). Libraries were sequenced (paired end, 2 x 50 bp) on an Illumina Nova seq 6000 SP100 with ∼50M reads/sample. Raw read files were uploaded to the European Nucleotide Archive and are available under accession PRJEB85227. Read quality was assessed by fastqc (v.0.11.8) prior to read alignment via HISAT2 (v.2.1.0) against the University of Liverpool *F. hepatica* genome [47] using genome and GTF files available on Wormbase ParaSite (WBPS19, https://parasite.wormbase.org/Fasciola_hepatica_prjeb58756/Info/Index/).

Transcripts were assembled in Stringtie (v1.3.6) and a master reference file, excluding isoforms, was generated by merging transcripts from different replicates with the reference genome. This master reference file was used to generate gene counts via Stringtie (v1.3.6) before export into a format readable by DESeq2. ‘Novel’ transcripts (MSTRGs) generated by Stringtie (v1.3.6) can be found in S2 File.

‘Novel’ MSTRG sequences with no BLAST hit in the NCBI nr database other than *F. hepatica* and genes with <10 counts across all samples were removed before differential gene expression analysis in R (v4.2.1) by DESeq2 (v.1.34.0). A p-value threshold of 0.001 false discovery rate (FDR) was set to identify differentially expressed genes (DEGs). These DEGs were annotated using the OmicsBox BLAST2GO suite [147] via BLASTx searches against the NCBI non-redundant, landmark and SwissProt databases. Additional BLASTs were carried out against the *S. mansoni* genome (v10, WBPS19, https://parasite.wormbase.org/Schistosoma_mansoni_prjea36577/Info/Index/). Gene ontology (GO) terms were mapped to genes based on BLAST searches and InterproScan results by BLAST2GO annotation software [147] before TOPGO (v.2.36) analysis (parameters: FDR <0.05, method = weight01, statistic = fisher) was carried out on DEGs. KEGG IDs were assigned to transcripts using the eggNOG-mapper [148] tool in OmicsBox (https://www.biobam.com/omicsbox) before gage (v.2.44) and pathview (v.1.34) were used to identify up and downregulated pathways. TOPGO (v.2.36) analysis of *S. mansoni* upregulated genes from Collins et al. [13] was performed by updating all genes to their current IDs and downloading GO terms from Wormbase ParaSite (WBPS19). Parameters were as those used for *F. hepatica* TOPGO analysis. R scripts are available on Github (https://github.com/pmccusker09/F.hepatica_plk1-RNAi_transcriptome_R_analysis.git).

Remaining total RNA was processed by the Genomics Core Technology Unit at Queen’s University Belfast for miRNA sequencing. Briefly, total RNA was library prepped with the QIASeq miRNA prep kit (#331502, Qiagen) and sequenced on an Illumina Next Seq 2000 with ∼8 M single reads per sample (100 bp). Adaptor sequences were trimmed using trimgalore (v.0.4.4) and read quality was assessed by fastqc (v.0.11.8). Raw read files are available under accession PRJEB85228 at the European Nucleotide Archive. Reads were mapped to the University of Liverpool genome downloaded from WormBase ParaSite (WBPS18, https://parasite.wormbase.org/Fasciola_hepatica_prjeb25283/Info/Index/) by bowtie2 (v.2.3.5.1) and miRDeep2 (v.0.1.3) before miRNA counts were generated by miRDeep2 (v.0.1.3), using predicted mature *F. hepatica* miRNAs [132] as a guide. Only counts that mapped to miRNAs previously identified by Herron et al. [132] were taken forward for differential expression analysis by DESeq2 (v.1.34.0) in R (v4.2.1). The targets of differentially expressed miRNAs were predicted by using the same thresholds as Gillan et al. [149]; miRanda (v.; total score >145, energy < -10), RNAhybrid (v.; p<0.1, energy < -22) and PITA (v.; seed sequence of 8 bases with DDG < -10). Target prediction was based on the WBPS18 version of the *F. hepatica* genome, as the miRNAs were predicted from that assembly [132], and so RNA-seq analysis was also carried out against this previous version of the genome assembly using the same methodology as detailed above.

### Exposure of juvenile *F. hepatica* to PLK1 inhibitor *in vitro*

The PLK inhibitor, BI 2536 (Millipore Sigma) was chosen for testing on juvenile *F. hepatica in vitro*. BI 2536 has a strong selectivity for PLK1 and so the amino acid sequence of FhPLK1 was examined to determine if key amino acid residues implicated in drug binding were conserved [150]. The PLK1 sequences of *F. hepatica*, *H. sapiens* [P53350], *S. mansoni* [Q5UES2] and *E. multilocularis* [U6HQM1], were aligned using Clustal Omega (https://www.ebi.ac.uk/Tools/msa/clustalo/). Other putative *F. hepatica* PLKs identified in our bioinformatic screen were included in the alignment to help predict BI 2536 activity. Further BLAST searches against the *F. hepatica* predicted proteomes (WBPS19, https://parasite.wormbase.org) with *H. sapiens* PLKs1-5 [P53350; Q9NYY3; Q9H4B4; O00444; Q496M5] as query sequences did not identify additional *F. hepatica* PLKs.

BI 2536 (#A10134, AdooQ Bioscience) and TCBZ (#1681611, Millipore Sigma) were dissolved in 1.7 mL NoStick hydrophobic microtubes (#1210-S0, SSIbio) using dimethyl sulphoxide (DMSO). Juvenile *F. hepatica* were washed three times in 1 mL RPMI 1640 to remove excess CS50. They were then transferred to 3.5 cm petri dishes (#82.1135.500, Sarstedt) for exposure in up to 10 µM test compound in a final volume of 3 mL RPMI 1640 for 18 h in standard culture conditions. Compounds were added such that the final DMSO concentration was 0.1% (v/v) in 3 mL RPMI. A vehicle control group (DMSO-treated) was included, with treatments carried out in triplicate. At the conclusion of drug incubations juveniles were subjected to phenotypic observation as detailed above. Where NEJs/juveniles were being further cultured post-exposure, they were washed three times in 1 mL RPMI 1640 to remove excess drug, prior to further culturing as previously described.

### Statistical Analyses

Unless otherwise stated graphs were generated and statistical analyses performed in GraphPad Prism (v10, La Jolla, CA USA). All datasets were tested for normality using either Kolmogorov-Smirnov or Shapiro-Wilk tests. Where data were normally distributed, parametric tests were performed, whereas nonparametric tests were employed where data were non-normally distributed. Relevant post-hoc tests (e.g. Dunn’s, Dunnett’s and Tukey’s multiple comparisons tests) were performed to identify significant differences between multiple groups. Volcano plots were produced using R (v4.2.1). Mean ±standard error mean (SEM) is displayed on graphs unless otherwise stated.

## Supplementary Information

**S1 Fig. Identification and compilation of *Fasciola hepatica* protein kinases.**

(A) Bioinformatics pipeline used to identify putative protein kinases in *F. hepatica* through HMM searches, BLAST identification, manual curation and CLANs analysis. (B) Proportions (%) of eukaryotic protein kinase families in *F. hepatica*, *Schistosoma mansoni*, *Schistosoma haematobium*, *Caenorhabditis elegans*, *Haemonchus contortus*, *Drosophila melanogaster* and *Homo sapiens* kinomes. (C) Maximum-likelihood tree (WAG +G + F model) with 100 bootstraps of predicted *F. hepatica* atypical protein kinases; colours denote family.

**S2 Fig. Cell proliferation does not recover up to a week after *fhplk1* dsRNA exposure in juvenile Fasciola hepatica in vitro.**

(A) Timelines for experiments show juvenile *F. hepatica* were treated for four weeks with either control dsRNA or *fhplk1* dsRNA before a final week in culture with no dsRNA treatments prior to EdU exposure and subsequent staining. (B) Confocal images of EdU staining (green) in juvenile *F. hepatica* treated according to adjacent timelines show that EdU^+^ nuclei did not recover after *fhplk1* dsRNA exposures were stopped; scale bar = 100 µm, DAPI counterstain (magenta). (C) Mean # EdU^+^ nuclei ±SEM in juvenile *F. hepatica* treated according to adjacent timelines shows complete loss of EdU^+^ nuclei in *fhplk1* dsRNA-treated worms, even after culture for one week post drug exposure.

**S3 Fig. Downregulation of cell cycle and ribosome associated transcripts in *fhplk1*-RNAi juvenile Fasciola hepatica.**

(A) Mean area (µm^2^) ±SEM of juvenile *F. hepatica* used in transcriptomics shows reduced growth rate in worms repeatedly treated with *fhplk1* dsRNA for three weeks. (B) Confocal images of EdU staining (green) in juvenile *F. hepatica* used for transcriptomics confirmed ablation of EdU^+^ nuclei after repeated treatment with *fhplk1* dsRNA for three weeks. (C) GO terms (top 10 molecular function and top 10 cellular component) overrepresented among downregulated transcripts following *fhplk1* dsRNA treatment in juvenile *F. hepatica*. (D) KEGG pathways significantly downregulated following *fhplk1*-RNAi in juvenile *F. hepatica* are associated with ribosomal components and cell cycle/proliferation. (E) Integrative genomics viewer (IGV) screenshot of transcriptomic reads in all RNA sequenced samples mapped against the *F. hepatica fhplk1* gene shows reads across the whole gene in control dsRNA samples, but a spike in reads around the dsRNA amplicon region in *fhplk1* dsRNA samples (red box to highlight). (F) IGV screenshots of beginning and end of *fhplk1* dsRNA amplicon region show that reads in *fhplk1* dsRNA samples begin and end where forward (‘F’ green) and reverse (‘R’ pink) amplicon primers are located.

**S4 Fig. Upregulated cell signalling and receptors/ion channel subunits in *fhplk1*-RNAi *Fasciola hepatica* juveniles.**

(A) GO terms (top 10 molecular function and top 10 cellular component) overrepresented in transcripts upregulated following *fhplk1*-RNAi in *F. hepatica* juveniles. (B) KEGG pathways associated with inter-cell signalling (gap junction, neural synapses and signalling pathways) significantly upregulated following *fhplk1*-RNAi in juvenile *F. hepatica*. (C) KEGG Gap Junction pathway following *fhplk1*-RNAi shows upregulation of diverse pathway components (red = upregulated; grey = no change; blue = downregulated; white = unassigned KEGG ID). (D) KEGG GABAergic synapse pathway following *fhplk1*-RNAi shows upregulation of diverse pathway components (red = upregulated; grey = no change; blue = downregulated; white = unassigned KEGG ID). (E) Top 10 Biological process GO terms overrepresented in significantly upregulated transcripts in both *fhplk1*-RNAi and slower growing *in vitro* worms (S6 Table) are associated with neuronal signalling. (F) DAPI (nuclei) stain coverage (as a percentage of the total area of the worm) in control-dsRNA treated and *fhplk1*-dsRNA treated worms shows ∼20% reduction. (G) Significantly upregulated receptors/channel subunits following *fhplk1*-RNAi.

**S5 Fig. GO terms of predicted targets for differentially expressed miRNAs following *fhplk1*-RNAi in *Fasciola hepatica* juveniles do not match the GO terms associated with differentially expressed mRNAs.**

(A) Volcano plot of miRNA expression (red dots indicate differentially expressed miRNA) in *fhplk1*-RNAi juvenile *F. hepatica*. (B) Mean Log_2_foldchange and frequency of GO terms for the predicted target transcripts of differentially expressed miRNAs following *fhplk1*-RNAi in *F. hepatica* juveniles.

**S6 Fig. The PLK1 inhibitor BI 2536 does not affect *Fasciola hepatica* motility *in vitro*.**

(A) Conservation of BI 2536 binding residues within *F. hepatica* FhPLK1, *Echinococcus multilocularis* EmPLK1, *Schistosoma mansoni* SmPLK1 and *Homo sapiens* PLK1 kinase domains, but not FhPLK2 or FhPLK4 (residues implicated in BI 2536 binding highlighted in black; Leu132 residue critical to drug interaction in *H. sapiens* PLK1 highlighted in red and marked with a red arrow). (B) Worm motility (%)

±SEM of *F. hepatica* newly excysted juveniles (NEJs) 18 hours after *in vitro* BI 2536 treatment. (C)

Motility profile of *F. hepatica* NEJs treated with 10 µM BI 2536 (blue), 0.1% DMSO (purple) and 1 µM TCBZ (green). Each line represents a single parasite; data presented as worm length (µm) over time (frames).

**S1 Table. Summary table of predicated *Fasciola hepatica* kinome based on bioinformatic screen.** Genes IDs, BLAST hits, IDs in alternative *F. hepatica* genome assembly listed. Eukaryotic protein kinases and atypical protein kinases in separate sheets.

**S2 Table. Summary of kinases downregulated following irradiation of *Fasciola hepatica* juveniles and upregulated in faster-growing *in vivo F. hepatica* juveniles.**

**S3 Table. DESeq2 results after three weeks of *fhplk1* RNAi in *Fasciola hepatica* juveniles *in vitro*.** (Sheet 1) Downregulated genes (adjp < 0.001), (Sheet 2) Upregulated genes (adjp < 0.001), (Sheet 3) All DESeq2 results.

**S4 Table. Common downregulated genes following *fhplk1* RNAi in *Fasciola hepatica* juveniles, irradiation in *F. hepatica* juveniles and irradiation in *Schistosoma mansoni* adults.**

(Sheet 1) Genes downregulated in both *fhplk1-*RNAi-*F. hepatica* juveniles, irradiated *F. hepatica* juveniles and irradiated *S. mansoni* adults. (Sheet 2) Genes downregulated in both irradiated *F. hepatica* juveniles and irradiated *S. mansoni* adults. (Sheet 3) Genes downregulated in both *fhplk1-*RNAi-*F. hepatica* juveniles and irradiated *S. mansoni* adults. (Sheet 4) Genes downregulated in both *fhplk1-* RNAi-*F. hepatica* juveniles and irradiated *F. hepatica* juveniles.

**S5 Table. Log_2_FoldChange of predicted *Fasciola hepatica* neuropeptides following RNAi of *fhplk1* in juveniles across three weeks *in vitro*.**

**S6 Table. Shared genes that are upregulated following *fhplk1* RNAi and downregulated in faster-growing *in vivo* juveniles.**

**S7 Table. Comparison of fold change for signalling-associated genes following *fhplk1* RNAi or in faster-growing *in vivo* juveniles.**

**S8 Table. GO terms upregulated in adult *Schistosoma mansoni* following neoblast-like cell ablation.** (Sheet 1) GO terms upregulated 14 days after irradiation. (Sheet 2) GO terms upregulated 21 days after *smh2b* RNAi. (Sheet 3) GO terms upregulated 21 days after *smfgfrA* RNAi.

**S9 Table. Differentially expressed miRNAs following *fhplk1* RNAi in *Fasciola hepatica* juveniles.** (Sheet 1) Differentially expressed miRNAs. (Sheet 2) Predicted targets of differentially expressed miRNAs.

**S10 Table. Primers used in PCR reactions.**

**S11 Table. Raw values used to produce graphs in Figs 1, 3, 4, 5, 6, S1, S2, S3, S4 and S6.**

**S1 File. Fasta file of all sequences in predicted *Fasciola hepatica* kinome**

**S2 File. Nucleotide sequences of ‘novel’ MSTRG *Fasciola hepatica* transcripts identified by Stringtie during transcriptomic processing.**

## References

1. FAO. Diseases in domestic animals caused by flukes. Food and Agriculture Organisation, Rome. Rome: Food and Agriculture Organisation of the United Nations; 1994.

2. WHO. Integrating neglected tropical diseases into global health and development: fourth WHO report on neglected tropical diseases. Geneva; 2017. Available: https://unitingtocombatntds.org/wp-content/uploads/2017/11/4th_who_ntd_report.pdf

3. Kelley JM, Elliott TP, Beddoe T, Anderson G, Skuce P, Spithill TW. Current threat of triclabendazole resistance in Fasciola hepatica. Trends Parasitol. 2016;32: 458–469. doi:10.1016/j.pt.2016.03.002

4. McVeigh P, McCusker P, Robb E, Wells D, Gardiner E, Mousley A, et al. Reasons to Be Nervous about Flukicide Discovery. Trends Parasitol. 2018;34: 184–196. doi:10.1016/j.pt.2017.11.010

5. Boray JC. Studies on experimental infections with fasciola hepatica, with particular reference to acute fascioliasis in sheep. Ann Trop Med Parasitol. 1967;61: 439–450. doi:10.1080/00034983.1967.11686513

6. Fairweather I, Boray JC. Fasciolicides: efficacy, actions, resistance and its management. The Veterinary Journal. 1999;158: 81–112. doi:10.1053/TVJL.1999.0377

7. Wagner DE, Wang IE, Reddien PW. Clonogenic neoblasts are pluripotent adult stem cells that underlie planarian regeneration. Science. 2011;332: 811–816. doi:10.1126/science.1203983

8. Wolff É, Dubois F. Sur la migration des cellules de régénération chez les Planaires. Revue Suisse De Zoologie. 1948;55: 218–227. doi:10.5962/bhl.part.117877

9. Campbell NJ, Gregg P, Kelly JD, Dineen JK. Failure to induce homologous immunity to Fasciola hepatica in sheep vaccinated with irradiated metacercariae. Vet Parasitol. 1978;4: 143–152. doi:10.1016/0304-4017(78)90005-5

10. Creaney J, Spithill TW, Thompson CM, Wilson LR, Sandeman RM, Parsons JC. Attempted immunisation of sheep against Fasciola hepatica using γ-irradiated metacercariae. Int J Parasitol. 1995;25: 853–856. doi:10.1016/0020-7519(94)00204-2

11. Wang B, Collins JJ, Newmark PA. Functional genomic characterization of neoblast-like stem cells in larval Schistosoma mansoni. Elife. 2013;2: e00768. doi:10.7554/eLife.00768

12. Collins III JJ, Wang B, Lambrus BG, Tharp ME, Iyer H, Newmark PA. Adult somatic stem cells in the human parasite Schistosoma mansoni. Nature. 2013;494: 476–9. doi:10.1038/nature11924

13. Collins III JJ, Wendt GR, Iyer H, Newmark PA. Stem cell progeny contribute to the schistosome host-parasite interface. Elife. 2016;5: e12473. doi:10.7554/eLife.12473

14. Wendt GR, Collins JN, Pei J, Pearson MS, Bennett HM, Loukas A, et al. Flatworm-specific transcriptional regulators promote the specification of tegumental progenitors in Schistosoma mansoni. eLife. 2018;7: e33221. doi:10.7554/eLife.33221

15. Koziol U, Rauschendorfer T, Zanon Rodríguez L, Krohne G, Brehm K. The unique stem cell system of the immortal larva of the human parasite Echinococcus multilocularis. Evodevo. 2014;5: 10. doi:10.1186/2041-9139-5-10

16. Rozario T, Quinn EB, Wang J, Davis RE, Newmark PA. Region-specific regulation of stem cell-driven regeneration in tapeworms. Elife. 2019;8: e48958. doi:10.7554/eLife.48958

17. Collins JNR, Collins III JJ. Tissue degeneration following loss of Schistosoma mansoni cbp1 is associated with increased stem cell proliferation and parasite death in vivo. PLoS Pathog. 2016;12: e1005963. doi:10.1371/journal.ppat.1005963

18. Wendt G, Shiroor DA, Adler CE, Collins III JJ. Convergent evolution of “genome guardian” functions in a parasite-specific p53 homolog. bioRxiv. 2021; 2021.11.28.470261. doi:10.1101/2021.11.28.470261

19. Romero AA, Cobb SA, Collins JNR, Kliewer SA, Mangelsdorf DJ, Collins III JJ. The Schistosoma mansoni nuclear receptor FTZ-F1 maintains esophageal gland function via transcriptional regulation of meg-8.3. PLoS Pathog. 2021;17: e1010140. doi:10.1371/journal.ppat.1010140

20. McCusker P, McVeigh P, Rathinasamy V, Toet H, McCammick E, O’Connor A, et al. Stimulating neoblast-like cell proliferation in juvenile Fasciola hepatica supports growth and progression towards the adult phenotype in vitro. PLoS Negl Trop Dis. 2016;10: e0004994. doi:10.1371/journal.pntd.0004994

21. McCusker P, Clarke NG, Erica G, Armstrong R, McCammick EM, McVeigh P, et al. Neoblast-like stem cells of Fasciola hepatica. PLoS Pathog. 2024;20: e1011903. doi:10.1371/journal.ppat.1011903

22. Fabbro D, Cowan-Jacob SW, Moebitz H. Ten things you should know about protein kinases: IUPHAR Review 14. Br J Pharmacol. 2015;172: 2675–2700. doi:10.1111/BPH.13096

23. Manning G, Whyte DB, Martinez R, Hunter T, Sudarsanam S. The protein kinase complement of the human genome. Science. 2002;298: 1912–1934. doi:10.1126/science.1075762

24. Sharma K, D’Souza RCJ, Tyanova S, Schaab C, Wiśniewski JR, Cox J, et al. Ultradeep Human Phosphoproteome Reveals a Distinct Regulatory Nature of Tyr and Ser/Thr-Based Signaling. Cell Rep. 2014;8: 1583–1594. doi:10.1016/j.celrep.2014.07.036

25. Wilson LJ, Linley A, Hammond DE, Hood FE, Coulson JM, MacEwan DJ, et al. New Perspectives, opportunities, and challenges in exploring the human protein kinome. Cancer Res. 2018;78: 15–29. doi:10.1158/0008-5472.can-17-2291

26. Roskoski R. Properties of FDA-approved small molecule protein kinase inhibitors. Pharmacol Res. 2019;144: 19–50. doi:10.1016/j.phrs.2019.03.006

27. Gramberg S, Puckelwaldt O, Schmitt T, Lu Z, Haeberlein S. Spatial transcriptomics of a parasitic flatworm provides a molecular map of drug targets and drug resistance genes. Nat Commun. 2024;15: 1–19. doi:10.1038/s41467-024-53215-3

28. Grevelding CG, Langner S, Dissous C. Kinases: Molecular Stage Directors for Schistosome Development and Differentiation. Trends Parasitol. 2018;34: 246–260. doi:10.1016/j.pt.2017.12.001

29. Arora N, Raj A, Anjum F, Kaur R, Rawat SS, Kumar R, et al. Unveiling Taenia solium kinome profile and its potential for new therapeutic targets. Expert Rev Proteomics. 2020;17: 85–94. doi:10.1080/14789450.2020.1719835

30. Stroehlein AJ, Young ND, Jex AR, Sternberg PW, Tan P, Boag PR, et al. Defining the Schistosoma haematobium kinome enables the prediction of essential kinases as anti-schistosome drug targets. Sci Rep. 2015;5. doi:10.1038/srep17759

31. Andrade LF, Nahum LA, Avelar LGA, Silva LL, Zerlotini A, Ruiz JC, et al. Eukaryotic Protein Kinases (ePKs) of the Helminth Parasite Schistosoma mansoni. BMC Genomics. 2011;12: 1–19. doi:10.1186/1471-2164-12-215

32. Sarkar N, Singh A, Kumar P, Kaushik M. Protein kinases: Role of their dysregulation in carcinogenesis, identification and inhibition. Drug Res. 2023;73: 189–199. doi:10.1055/A-1989-1856

33. Llamazares S, Moreira A, Tavares A, Girdham C, Spruce BA, Gonzalez C, et al. polo encodes a protein kinase homolog required for mitosis in Drosophila. Genes Dev. 1991;5: 2153–2165. doi:10.1101/gad.5.12A.2153

34. Sunkel CE, Glover DM. Polo, a mitotic mutant of Drosophila displaying abnormal spindle poles. J Cell Sci. 1988;89: 25–38. doi:10.1242/jcs.89.1.25

35. Kitada K, Johnson AL, Johnston LH, Sugino A. A Multicopy Suppressor Gene of the Saccharomyces cerevisiae G1 Cell Cycle Mutant Gene dbf4 Encodes a Protein Kinase and Is Identified as CDC5. Mol Cell Biol. 1993;13: 4445–4457. doi:10.1128/MCB.13.7.4445-4457.1993

36. Ohkura H, Hagan IM, Glover DM. The conserved Schizosaccharomyces pombe kinase plo1, required to form a bipolar spindle, the actin ring, and septum, can drive septum formation in G1 and G2 cells. Genes Dev. 1995;9: 1059–1073. doi:10.1101/GAD.9.9.1059

37. Hamanaka R, Maloid S, Smith MR, O CD, Ferris DK, Li DL, et al. Cloning and characterization of human and murine homologues of the Drosophila polo serine-threonine kinase. Cell Growth & Differ. 1994;5: 249–257.

38. Zitouni S, Nabais C, Jana SC, Guerrero A, Bettencourt-Dias M. Polo-like kinases: structural variations lead to multiple functions. Nat Rev Mol Cell Biol. 2014;15: 433–452. doi:10.1038/nrm3819

39. Iliaki S, Beyaert R, Afonina IS. Polo-like kinase 1 (PLK1) signaling in cancer and beyond. Biochem Pharmacol. 2021;193: 114747. doi:10.1016/j.bcp.2021.114747

40. Long T, Cailliau K, Beckmann S, Browaeys E, Trolet J, Grevelding CG, et al. Schistosoma mansoni Polo-like kinase 1: A mitotic kinase with key functions in parasite reproduction. Int J Parasitol. 2010;40: 1075–1086. doi:10.1016/j.ijpara.2010.03.002

41. Dissous C, Grevelding CG, Long T. Schistosoma mansoni polo-like kinases and their function in control of mitosis and parasite reproduction. An Acad Bras Cienc. 2011;83: 627–635. doi:10.1590/S0001-37652011000200022

42. Long T, Neitz RJ, Beasley R, Kalyanaraman C, Suzuki BM, Jacobson MP, et al. Structure-Bioactivity Relationship for Benzimidazole Thiophene Inhibitors of Polo-Like Kinase 1 (PLK1), a Potential Drug Target in Schistosoma mansoni. PLoS Negl Trop Dis. 2016;10: e0004356. doi:10.1371/journal.pntd.0004356

43. Schubert A, Koziol U, Cailliau K, Vanderstraete M, Dissous C, Brehm K. Targeting Echinococcus multilocularis stem cells by inhibition of the polo-like kinase Emplk1. PLoS Negl Trop Dis. 2014;8: e2870. doi:10.1371/journal.pntd.0002870

44. Houhou H, Puckelwaldt O, Strube C, Haeberlein S. Reference gene analysis and its use for kinase expression profiling in Fasciola hepatica. Sci Rep. 2019;9: 15867. doi:10.1038/s41598-019-52416-x

45. Morawietz CM, Houhou H, Puckelwaldt O, Hehr L, Dreisbach D, Mokosch A, et al. Targeting kinases in Fasciola hepatica: anthelminthic effects and tissue distribution of selected kinase inhibitors. Front Vet Sci. 2020;7: 611270. doi:10.3389/fvets.2020.611270

46. Robb E, McCammick EM, Wells D, McVeigh P, Gardiner E, Armstrong R, et al. Transcriptomic analysis supports a role for the nervous system in regulating growth and development of Fasciola hepatica juveniles. PLoS Negl Trop Dis. 2022;16: e0010854. doi:10.1371/journal.pntd.0010854

47. Cwiklinski K, Dalton JP, Dufresne PJ, La Course J, Williams DJ, Hodgkinson J, et al. The Fasciola hepatica genome: gene duplication and polymorphism reveals adaptation to the host environment and the capacity for rapid evolution. Genome Biol. 2015;16: 71. doi:10.1186/s13059-015-0632-2

48. Plowman GD, Sudarsanam S, Bingham J, Whyte D, Hunter T. The protein kinases of Caenorhabditis elegans: A model for signal transduction in multicellular organisms. Proc Natl Acad Sci USA. 1999;96: 13603–13610. doi:10.1073/pnas.96.24.13603

49. Manning G, Plowman GD, Hunter T, Sudarsanam S. Evolution of protein kinase signaling from yeast to man. Trends Biochem Sci. 2002;27: 514–520. doi:10.1016/S0968-0004(02)02179-5

50. Skelding KA, Rostas JAP. The Role of Molecular Regulation and Targeting in Regulating Calcium/Calmodulin Stimulated Protein Kinases. Adv Exp Med Biol. 2012;740: 703–730. doi:10.1007/978-94-007-2888-2_31

51. You H, McManus DP, Hu W, Smout MJ, Brindley PJ, Gobert GN. Transcriptional Responses of In Vivo Praziquantel Exposure in Schistosomes Identifies a Functional Role for Calcium Signalling Pathway Member CamKII. PLoS Pathog. 2013;9: e1003254. doi:10.1371/journal.ppat.1003254

52. Hirst NL, Lawton SP, Walker AJ. CaMKII regulates neuromuscular activity and survival of the human blood fluke Schistosoma mansoni. Sci Rep. 2022;12: 1–14. doi:10.1038/s41598-022-23962-8

53. Chowdhury I, Dashi G, Keskitalo S. CMGC Kinases in Health and Cancer. Cancers 2023, Vol 15, Page 3838. 2023;15: 3838. doi:10.3390/cancers15153838

54. Puckelwaldt O, Gramberg S, Ajmera S, Koepke J, Samakovlis C, Haeberlein S. Single-cell transcriptomics identifies a p21-activated kinase important for survival of the zoonotic parasite Fasciola hepatica. bioRxiv. 2024; 2024.03.26.586785. doi:10.1101/2024.03.26.586785

55. Kanev GK, de Graaf C, de Esch IJP, Leurs R, Würdinger T, Westerman BA, et al. The Landscape of Atypical and Eukaryotic Protein Kinases. Trends Pharmacol Sci. 2019;40: 818–832. doi:10.1016/j.tips.2019.09.002

56. Zaru R, Magrane M, O’Donovan C, Bateman A, Martin MJ, Alpi E, et al. From the research laboratory to the database: the Caenorhabditis elegans kinome in UniProtKB. Biochemical Journal. 2017;474: 493–515. doi:10.1042/BCJ20160991

57. Hua H, Kong Q, Zhang H, Wang J, Luo T, Jiang Y. Targeting mTOR for cancer therapy. J Hematol Oncol. 2019;12: 1–19. doi:10.1186/S13045-019-0754-1

58. Weber AM, Ryan AJ. ATM and ATR as therapeutic targets in cancer. Pharmacol Ther. 2015;149: 124–138. doi:10.1016/j.pharmthera.2014.12.001

59. Burke JE, Triscott J, Emerling BM, Hammond GRV. Beyond PI3Ks: targeting phosphoinositide kinases in disease. Nature Reviews Drug Discovery 2022 22:5. 2022;22: 357–386. doi:10.1038/s41573-022-00582-5

60. Malumbres M, Barbacid M. Cell cycle, CDKs and cancer: a changing paradigm. Nat Rev Cancer. 2009;9: 153–166. doi:10.1038/nrc2602

61. Wang XQ, Lo CM, Chen L, Ngan ESW, Xu A, Poon RYC. CDK1-PDK1-PI3K/Akt signaling pathway regulates embryonic and induced pluripotency. Cell Death Differ. 2017;24: 38–48. doi:10.1038/cdd.2016.84

62. Santamaría D, Barrière C, Cerqueira A, Hunt S, Tardy C, Newton K, et al. Cdk1 is sufficient to drive the mammalian cell cycle. Nature 2007;448: 811–815. doi:10.1038/nature06046

63. Babina IS, Turner NC. Advances and challenges in targeting FGFR signalling in cancer. Nat Rev Cancer. 2017;17: 318–332. doi:10.1038/nrc.2017.8

64. Förster S, Koziol U, Schäfer T, Duvoisin R, Cailliau K, Vanderstraete M, et al. The role of fibroblast growth factor signalling in Echinococcus multilocularis development and host-parasite interaction. PLoS Negl Trop Dis. 2019;13: e0006959. doi:10.1371/journal.pntd.0006959

65. Du X, McManus DP, French JD, Collinson N, Sivakumaran H, MacGregor SR, et al. CRISPR interference for sequence-specific regulation of fibroblast growth factor receptor A in Schistosoma mansoni. Front Immunol. 2023;13: 1105719. doi:10.3389/fimmu.2022.1105719/bibtex

66. Roskoski R. Properties of FDA-approved small molecule protein kinase inhibitors: A 2021 update. Pharmacol Res. 2021;165: 105463. doi:10.1016/j.phrs.2021.105463

67. Strebhardt K, Ullrich A. Targeting polo-like kinase 1 for cancer therapy. Nat Rev Cancer. 2006;6: 321–330. doi:10.1038/nrc1841

68. Hanks SK, Hunter T. The eukaryotic protein kinase superfamily: kinase (catalytic) domain structure and classification1. The FASEB Journal. 1995;9: 576–596. doi:10.1096/fasebj.9.8.7768349

69. Seki A, Coppinger JA, Jang CY, Yates JR, Fang G. Bora and the kinase Aurora A cooperatively activate the kinase Plk1 and control mitotic entry. Science. 2008;320: 1655–1658. doi:10.1126/science.1157425

70. Elia AEH, Cantley LC, Yaffe MB. Proteomic screen finds pSer/pThr-binding domain localizing Plk1 to mitotic substrates. Science. 2003;299: 1228–1231. doi:10.1126/science.1079079

71. Elia AEH, Rellos P, Haire LF, Chao JW, Ivins FJ, Hoepker K, et al. The molecular basis for phosphodependent substrate targeting and regulation of Plks by the Polo-box domain. Cell. 2003;115: 83–95. doi:10.1016/S0092-8674(03)00725-6

72. Wang J, Chen R, Collins III JJ. Systematically improved in vitro culture conditions reveal new insights into the reproductive biology of the human parasite Schistosoma mansoni. PLoS Biol. 2019;17: e3000254. doi:10.1371/journal.pbio.3000254

73. Hanna R. Fasciola hepatica: Histology of the Reproductive Organs and Differential Effects of Triclabendazole on Drug-Sensitive and Drug-Resistant Fluke Isolates and on Flukes from Selected Field Cases. Pathogens. 2015;4: 431–456. doi:10.3390/pathogens4030431

74. Björkman N, Thorsell W. On the ultrastructure of the ovary of the liver fluke (Fasciola hepatica L.). Zeitschrift für Zellforschung und Mikroskopische Anatomie. 1964;63: 538–549. doi:10.1007/bf00339489/metrics

75. Marzluff WF, Wagner EJ, Duronio RJ. Metabolism and regulation of canonical histone mRNAs: life without a poly(A) tail. Nat Rev Genet. 2008;9: 843–854. doi:10.1038/nrg2438

76. Pettitt J, Crombie C, Schümperli D, Müller B. The Caenorhabditis elegans histone hairpin-binding protein is required for core histone gene expression and is essential for embryonic and postembryonic cell division. J Cell Sci. 2002;115: 857–866. doi:10.1242/JCS.115.4.857

77. Sullivan E, Santiago C, Parker ED, Dominski Z, Yang X, Lanzotti DJ, et al. Drosophila stem loop binding protein coordinates accumulation of mature histone mRNA with cell cycle progression. Genes Dev. 2001;15: 173–187. doi:10.1101/gad.862801

78. Solana J, Kao D, Mihaylova Y, Jaber-Hijazi F, Malla S, Wilson R, et al. Defining the molecular profile of planarian pluripotent stem cells using a combinatorial RNAseq, RNA interference and irradiation approach. Genome Biol. 2012;13: 1–23. doi:10.1186/gb-2012-13-3-r19

79. Spänkuch-Schmitt B, Bereiter-Hahn J, Kaufmann M, Strebhardt K. Effect of RNA Silencing of Polo-Like Kinase-1 (PLK1) on Apoptosis and Spindle Formation in Human Cancer Cells. J Natl Cancer Inst. 2002;94: 1863–1877. doi:10.1093/JNCI/94.24.1863

80. McCarroll JA, Dwarte T, Baigude H, Dang J, Yang L, Erlich RB, et al. Therapeutic targeting of polo-like kinase 1 using RNA-interfering nanoparticles (iNOPs) for the treatment of non-small cell lung cancer. Oncotarget. 2015;6: 12020–12034. doi:10.18632/oncotarget.2664

81. Sampath P, Pritchard DK, Pabon L, Reinecke H, Schwartz SM, Morris DR, et al. A Hierarchical Network Controls Protein Translation during Murine Embryonic Stem Cell Self-Renewal and Differentiation. Cell Stem Cell. 2008;2: 448–460. doi:10.1016/j.stem.2008.03.013

82. Saba JA, Liakath-Ali K, Green R, Watt FM. Translational control of stem cell function. Nat Rev Mol Cell Biol. 2021;22: 671–690. doi:10.1038/s41580-021-00386-2

83. Larson DE, Xie W, Glibetic M, O’Mahony D, Sells BH, Rothblum LI. Coordinated decreases in rRNA gene transcription factors and rRNA synthesis during muscle cell differentiation. PNAS. 1993;90: 7933–7936. doi:10.1073/PNAS.90.17.7933

84. Cong Y, Yang H, Zhang P, Xie Y, Cao X, Zhang L. Transcriptome Analysis of the Nematode Caenorhabditis elegans in Acidic Stress Environments. Front Physiol. 2020;11: 1107. doi:10.3389/fphys.2020.01107/full

85. Jasmer DP, Rosa BA, Mitreva M. Cell death and transcriptional responses induced in larvae of the nematode Haemonchus contortus by toxins/toxicants with broad phylogenetic efficacy. Pharmaceuticals. 2021;14: 598. doi:10.3390/ph14070598

86. Chen J. The Cell-Cycle Arrest and Apoptotic Functions of p53 in Tumor Initiation and Progression. Cold Spring Harb Perspect Med. 2016;6: a026104. doi:10.1101/cshperspect.A026104

87. Duffy MJ, Synnott NC, Crown J. Mutant p53 as a target for cancer treatment. Eur J Cancer. 2017;83: 258–265. doi:10.1016/J.EJCA.2017.06.023

88. Fei L, Xu H. Role of MCM2-7 protein phosphorylation in human cancer cells. Cell Biosci. 2018;8: 1–8. doi:10.1186/S13578-018-0242-2

89. McCusker P, Hussain W, McVeigh P, McCammick E, Clarke NG, Robb E, et al. RNA interference dynamics in juvenile Fasciola hepatica are altered during in vitro growth and development. Int J Parasitol Drugs Drug Resist. 2020;14: 46–55. doi:10.1016/j.ijpddr.2020.08.004

90. Vigneron S, Brioudes E, Burgess A, Labbé JC, Lorca T, Castro A. Greatwall maintains mitosis through regulation of PP2A. EMBO Journal. 2009;28: 2786–2793. doi:10.1038/emboj.2009.228

91. Im JS, Ki SH, Farina A, Jung DS, Hurwitz J, Lee JK. Assembly of the Cdc45-Mcm2-7-GINS complex in human cells requires the Ctf4/And-1, RecQL4, and Mcm10 proteins. PNAS. 2009;106: 15628–15632. doi:10.1073/pnas.0908039106

92. Komorowska K, Doyle A, Wahlestedt M, Subramaniam A, Debnath S, Chen J, et al. Hepatic Leukemia Factor Maintains Quiescence of Hematopoietic Stem Cells and Protects the Stem Cell Pool during Regeneration. Cell Rep. 2017;21: 3514–3523. doi:10.1016/j.celrep.2017.11.084

93. Stoll K, Bergmann M, Spiliotis M, Brehm K. A MEKK1 – JNK mitogen activated kinase (MAPK) cascade module is active in Echinococcus multilocularis stem cells. PLoS Negl Trop Dis. 2021;15: e0010027. doi:10.1371/journal.pntd.0010027

94. Beyer EC, Berthoud VM. Gap junction gene and protein families: Connexins, innexins, and pannexins. Biochim Biophys Acta Biomembr. 2018;1860: 5–8. doi:10.1016/j.bbamem.2017.05.016

95. Nogi T, Levin M. Characterization of innexin gene expression and functional roles of gap-junctional communication in planarian regeneration. Dev Biol. 2005;287: 314–335. doi:10.1016/j.ydbio.2005.09.002

96. Oviedo NJ, Morokuma J, Walentek P, Kema IP, Gu MB, Ahn JM, et al. Long-range neural and gap junction protein-mediated cues control polarity during planarian regeneration. Dev Biol. 2010;339: 188–199. doi:10.1016/j.ydbio.2009.12.012

97. Oviedo NJ, Levin M. smedinx-11 is a planarian stem cell gap junction gene required for regeneration and homeostasis. Development. 2007;134: 3121–3131. doi:10.1242/DEV.006635

98. Maule AG, Marks NJ, Day TA. Signalling Molecules and Nerve-Muscle Function. In: Maule AG, Marks N, editors. Parasitic Flatworms: Molecular Biology, Biochemistry, Immunology and Physiology. CABI; 2006. pp. 369–381.

99. El-Shehabi F, Ribeiro P. Histamine signalling in Schistosoma mansoni: Immunolocalisation and characterisation of a new histamine-responsive receptor (SmGPR-2). Int J Parasitol. 2010;40: 1395– 1406. doi:10.1016/J.ijpara.2010.04.006

100. Sukhodeo SC, Sangster NC, Mettrick DF. Effects of cholinergic drugs on longitudinal muscle contractions of Fasciola hepatica. J Parasitol. 1986;72: 858–864. doi:10.2307/3281834

101. Sukhdeo MVK, Mettrick DF. The behavior of juvenile Fasciola hepatica. J Parasitol. 1986;72: 492–497. doi:10.2307/3281496

102. Fanburg BL, Lee SL. A new role for an old molecule: serotonin as a mitogen. Am J Physiol. 1997;272: L795–806. doi:10.1152/ajplung.1997.272.5.L795

103. Franquinet R, Martelly I. Effects of serotonin and catecholamines on RNA synthesis in planarians; in vitro and in vivo studies. Cell Differ. 1981;10: 201–209. doi:10.1016/0045-6039(81)90002-6

104. Taft AS, Norante FA, Yoshino TP. The identification of inhibitors of Schistosoma mansoni miracidial transformation by incorporating a medium-throughput small-molecule screen. Exp Parasitol. 2010;125: 84–94. doi:10.1016/j.exppara.2009.12.021

105. Kawamoto F, Shozawa A, Kumada N, Kojima K. Possible roles of cAMP and Ca2+ in the regulation of miracidial transformation in Schistosoma mansoni. Parasitol Res. 1989;75: 368–374. doi:10.1007/BF00931132

106. Boyle JP, Zaide J V., Yoshino TP. Schistosoma mansoni: Effects of Serotonin and Serotonin Receptor Antagonists on Motility and Length of Primary Sporocysts in vitro. Exp Parasitol. 2000;94: 217–226. doi:10.1006/expr.2000.4500

107. Herz M, Brehm K. Serotonin stimulates Echinococcus multilocularis larval development. Parasit Vectors. 2021;14: 1–12. doi:10.1186/S13071-020-04533-0

108. Alqahtani S, Butcher MC, Ramage G, Dalby MJ, McLean W, Nile CJ. Acetylcholine Receptors in Mesenchymal Stem Cells. Stem Cells Dev. 2023;32: 47–59. doi:10.1089/SCD.2022.0201

109. Basu S, Dasgupta PS. Role of dopamine in malignant tumor growth. Endocrine. 2000;12: 237–241. doi:10.1385/endo:12:3:237

110. Lee I, Knickerbocker AC, Depew CR, Martin E, Dicent J, Miller GW, et al. Effect of altered production and storage of dopamine on development and behavior in C. elegans. bioRxiv. 2023; 2023.10.07.561350. doi:10.1101/2023.10.07.561350

111. Neckameyer WS. Multiple Roles for Dopamine in Drosophila Development. Dev Biol. 1996;176: 209–219. doi:10.1006/dbio.1996.0128

112. Taft AS, Norante FA, Yoshino TP. The identification of inhibitors of Schistosoma mansoni miracidial transformation by incorporating a medium-throughput small-molecule screen. Exp Parasitol. 2010;125: 84. doi:10.1016/j.exppara.2009.12.021

113. Rand JB, Russell RL. Choline acetyltransferase-deficient mutants of the nematode Caenorhabditis elegans. Genetics. 1984;106: 227–248. doi:10.1093/genetics/106.2.227

114. Zachary I, Woll PJ, Rozengurt E. A role for neuropeptides in the control of cell proliferation. Dev Biol. 1987;124: 295–308. doi:10.1016/0012-1606(87)90483-0

115. Kreshchenko ND. Functions of flatworm neuropeptides NPF, GYIRF and FMRF in course of pharyngeal regeneration of anterior body fragments of planarian, Girardia tigrina. Acta Biol Hung. 2008;59: 199–207. doi:10.1556/ABiol.59.2008.Suppl.29

116. Collins III JJ, Hou X, Romanova E V., Lambrus BG, Miller CM, Saberi A, et al. Genome-Wide Analyses Reveal a Role for Peptide Hormones in Planarian Germline Development. PLoS Biol. 2010;8: e1000509. doi:10.1371/journal.pbio.1000509

117. Saberi A, Jamal A, Beets I, Schoofs L, Newmark PA. GPCRs Direct Germline Development and Somatic Gonad Function in Planarians. PLoS Biol. 2016;14: e1002457. doi:10.1371/journal.pbio.1002457

118. Bagurija’ J, Salo E, Romero R. Effects of activators and antagonists of the neuropeptides substance P and substance K on cell proliferation in planarians. Int J Dev Biol. 1989;33: 261–264.

119. Li X, Weth O, Haimann M, Möscheid MF, Huber TS, Grevelding CG, et al. Rhodopsin orphan GPCR20 interacts with neuropeptides and directs growth, sexual differentiation, and egg production in female Schistosoma mansoni. Microbiol Spectr. 2023;12: e0219323 doi:10.1128/spectrum.02193-23

120. Almuedo-Castillo M, Sureda-Gómez M, Adell T. Wnt signaling in planarians: New answers to old questions. International Journal of Developmental Biology. 2012;56: 53–65. doi:10.1387/ijdb.113451ma

121. Ye Z, Xu J, Feng X, Jia Y, Fu Z, Hong Y, et al. Spatiotemporal expression pattern of Sjfz7 and its expression comparison with other frizzled family genes in developmental stages of Schistosoma japonicum. Gene Expression Patterns. 2019;32: 44–52. doi:10.1016/j.gep.2019.02.005

122. Cheng G, Li X, Qin F, Xu R, Zhang Y, Liu J, et al. Functional analysis of the Frzb2 gene in Schistosoma japonicum. Vet Res. 2019;50: 1–11. doi:10.1186/S13567-019-0716-1

123. Koziol U, Jarero F, Olson PD, Brehm K. Comparative analysis of Wnt expression identifies a highly conserved developmental transition in flatworms. BMC Biol. 2016;14: 1–16. doi:10.1186/s12915-016-0233-x

124. Gurley KA, Elliott SA, Simakov O, Schmidt HA, Holstein TW, Alvarado AS. Expression of secreted Wnt pathway components reveals unexpected complexity of the planarian amputation response. Dev Biol. 2010;347: 24–39. doi:10.1016/j.ydbio.2010.08.007

125. Adell T, Salò E, Boutos M, Bartscherer K. Smed-Evi/Wntless is required for β-catenin-dependent and-independent processes during planarian regeneration. Development. 2009;136: 905– 910. doi:10.1242/DEV.033761

126. Li HF, Wang XB, Jin YP, Xia YX, Feng XG, Yang JM, et al. Wnt4, the first member of the Wnt family identified in Schistosoma japonicum, regulates worm development by the canonical pathway. Parasitol Res. 2010;107: 795–805. doi:10.1007/S00436-010-1933-8

127. Kobayashi C, Saito Y, Ogawa K, Agata K. Wnt signaling is required for antero-posterior patterning of the planarian brain. Dev Biol. 2007;306: 714–724. doi:10.1016/j.ydbio.2007.04.010

128. Armstrong R, Marks NJ, Geary TG, Harrington J, Selzer PM, Maule AG. Wnt/β-catenin signalling underpins juvenile Fasciola hepatica growth and development. PLoS Pathog. 2025;21: e1012562. doi: 10.1371/journal.ppat.1012562

129. Lingueglia E, Champigny G, Lazdunski M, Barbry P. Cloning of the amiloride-sensitive FMRFamide peptide-gated sodium channel. Nature 1995;378: 730–733. doi:10.1038/378730a0

130. Baguñà J. The planarian neoblast: the rambling history of its origin and some current black boxes. Int J Dev Biol. 2012;56: 19–37. doi:10.1387/ijdb.113463jb

131. Yang D, Zhou Q, Labroska V, Qin S, Darbalaei S, Wu Y, et al. G protein-coupled receptors: structure- and function-based drug discovery. Signal Transduct Target Ther. 2021;6: 1–27. doi:10.1038/s41392-020-00435-w

132. Herron CM, O’Connor A, Robb E, McCammick E, Hill C, Marks NJ, et al. Developmental Regulation and Functional Prediction of microRNAs in an Expanded Fasciola hepatica miRNome. Front Cell Infect Microbiol. 2022;12. doi:10.3389/fcimb.2022.811123

133. Kothe M, Kohls D, Low S, Coli R, Rennie GR, Feru F, et al. Selectivity-determining Residues in Plk1. Chem Biol Drug Des. 2007;70: 540–546. doi:10.1111/J.1747-0285.2007.00594.X

134. Steegmaier M, Hoffmann M, Baum A, Lénárt P, Petronczki M, Krššák M, et al. BI 2536, a Potent and Selective Inhibitor of Polo-like Kinase 1, Inhibits Tumor Growth In Vivo. Curr Biol. 2007;17: 316– 322. doi:10.1016/j.cub.2006.12.037

135. Long T, Vanderstraete M, Cailliau K, Morel M, Lescuyer A, Gouignard N, et al. SmSak, the second polo-like kinase of the helminth parasite Schistosoma mansoni: conserved and unexpected roles in meiosis. PLoS One. 2012;7: e40045. doi:10.1371/journal.pone.0040045

136. Bennett CEE. Surface Features, Sensory Structures, and Movement of the Newly Excysted Juvenile Fasciola hepatica L. J Parasitol. 1975;61: 886–891.

137. Taylor S, Peters JM. Polo and Aurora kinases — lessons derived from chemical biology. Curr Opin Cell Biol. 2008;20: 77–84. doi:10.1016/J.CEB.2007.11.008

138. Gelmedin V, Dissous C, Grevelding CG. Re-positioning protein-kinase inhibitors against schistosomiasis. Future Med Chem. 2015;7: 737–752. doi:10.4155/fmc.15.31

139. McNulty SN, Tort JF, Rinaldi G, Fischer K, Rosa BA, Smircich P, et al. Genomes of Fasciola hepatica from the Americas Reveal Colonization with Neorickettsia Endobacteria Related to the Agents of Potomac Horse and Human Sennetsu Fevers. PLoS Genet. 2017;13: e1006537. doi:10.1371/journal.pgen.1006537

140. Lee RYN, Howe KL, Harris TW, Arnaboldi V, Cain S, Chan J, et al. WormBase 2017: molting into a new stage. Nucleic Acids Res. 2018;46: D869–D874. doi:10.1093/nar/gkx998

141. Tancred Frickey, Andrei Lupas. CLANS: a Java application for visualizing protein families based on pairwise similarity. Bioinformatics. 2004;20: 3702–3704. doi:10.1093/bioinformatics/BTH444

142. Tamura K, Stecher G, Kumar S. MEGA11: Molecular Evolutionary Genetics Analysis Version 11. Mol Biol Evol. 2021;38: 3022–3027. doi:10.1093/molbev/msab120

143. McVeigh P, McCammick EM, McCusker P, Morphew RM, Mousley A, Abidi A, et al. RNAi dynamics in juvenile Fasciola spp. liver flukes reveals the persistence of gene silencing in vitro. PLoS Negl Trop Dis. 2014;8: e3185. doi:10.1371/journal.pntd.0003185

144. Untergasser A, Nijveen H, Rao X, Bisseling T, Geurts R, Leunissen JAM. Primer3Plus, an enhanced web interface to Primer3. Nucleic Acids Res. 2007;35: W71–4. doi:10.1093/nar/gkm306

145. Schneider CA, Rasband WS, Eliceiri KW. NIH Image to ImageJ: 25 years of image analysis. Nat Methods 2012 9:7. 2012;9: 671–675. doi:10.1038/nmeth.2089

146. Pfaffl MW. A new mathematical model for relative quantification in real-time RT-PCR. Nucleic Acids Res. 2001;29: e45. doi:10.1093/nar/29.9.e45

147. Götz S, García-Gómez JM, Terol J, Williams TD, Nagaraj SH, Nueda MJ, et al. High-throughput functional annotation and data mining with the Blast2GO suite. Nucleic Acids Res. 2008;36: 3420–3435. doi:10.1093/nar/gkn176

148. Cantalapiedra CP, Hern̗andez-Plaza A, Letunic I, Bork P, Huerta-Cepas J. eggNOG-mapper v2: Functional Annotation, Orthology Assignments, and Domain Prediction at the Metagenomic Scale. Mol Biol Evol. 2021;38: 5825–5829. doi:10.1093/molbev/msab293

149. Gillan V, Maitland K, Laing R, Gu H, Marks ND, Winter AD, et al. Increased expression of a microRNA correlates with anthelmintic resistance in parasitic nematodes. Front Cell Infect Microbiol. 2017;7: 292984. doi:10.3389/FCIMB.2017.00452

150. Kothe M, Kohls D, Low S, Coli R, Rennie GR, Feru F, et al. Selectivity-determining Residues in Plk1. Chem Biol Drug Des. 2007;70: 540–546. doi:10.1111/J.1747-0285.2007.00594.X

