## Supplementary figures and images for "Polo-like kinase 1 regulates growth in juvenile *Fasciola hepatica*"

### S1_Fig

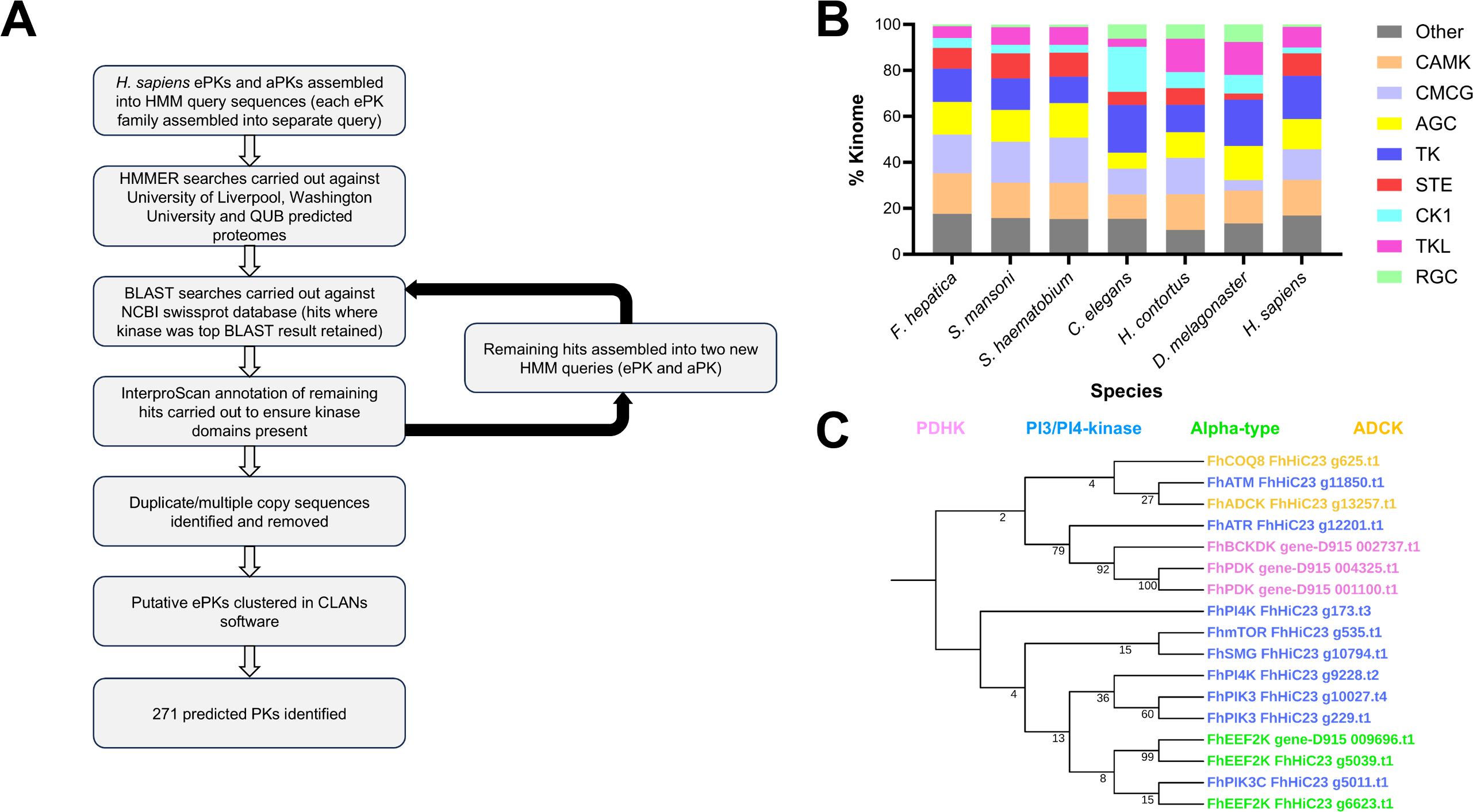

### S2_Fig

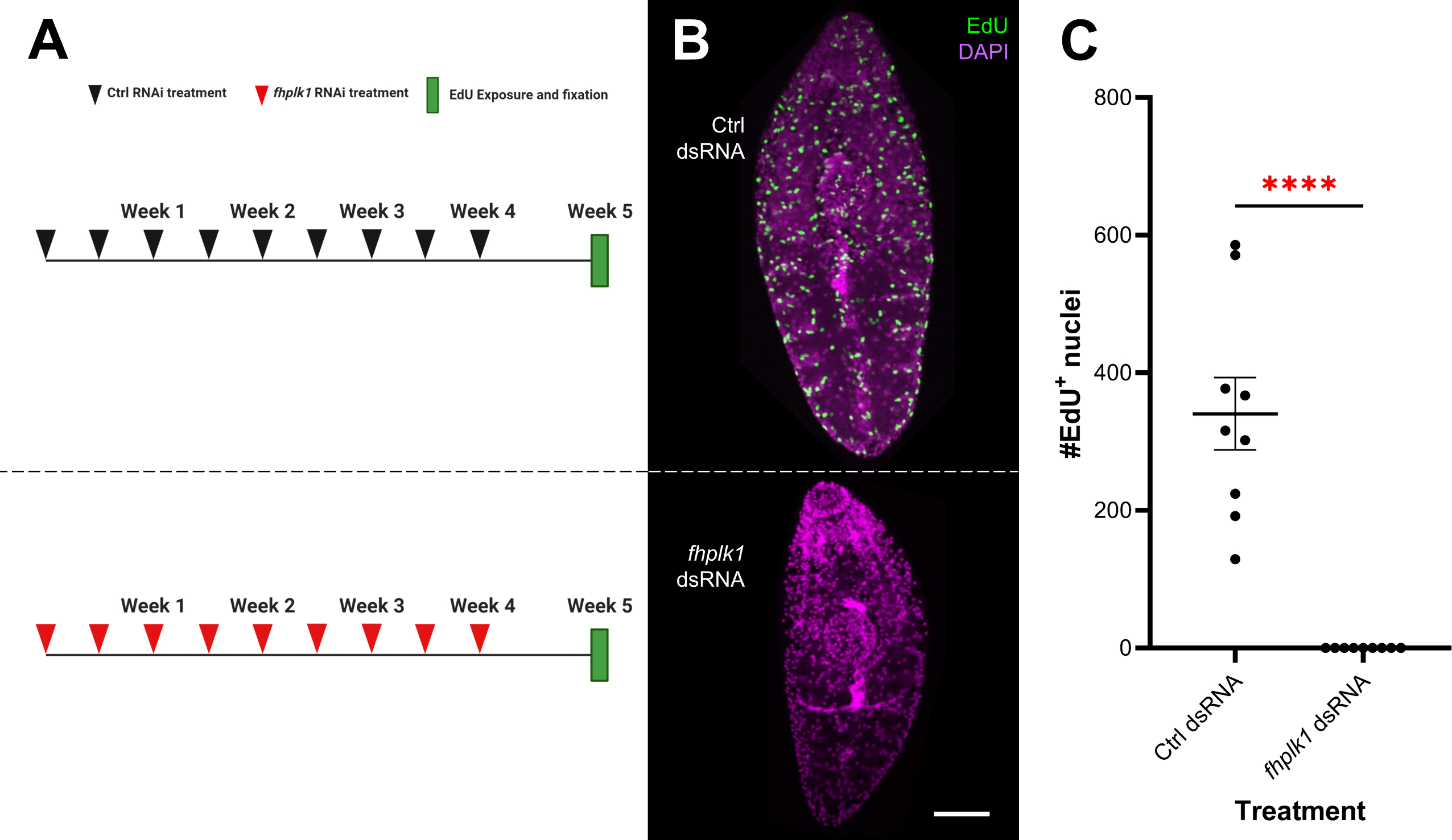

### S3_Fig

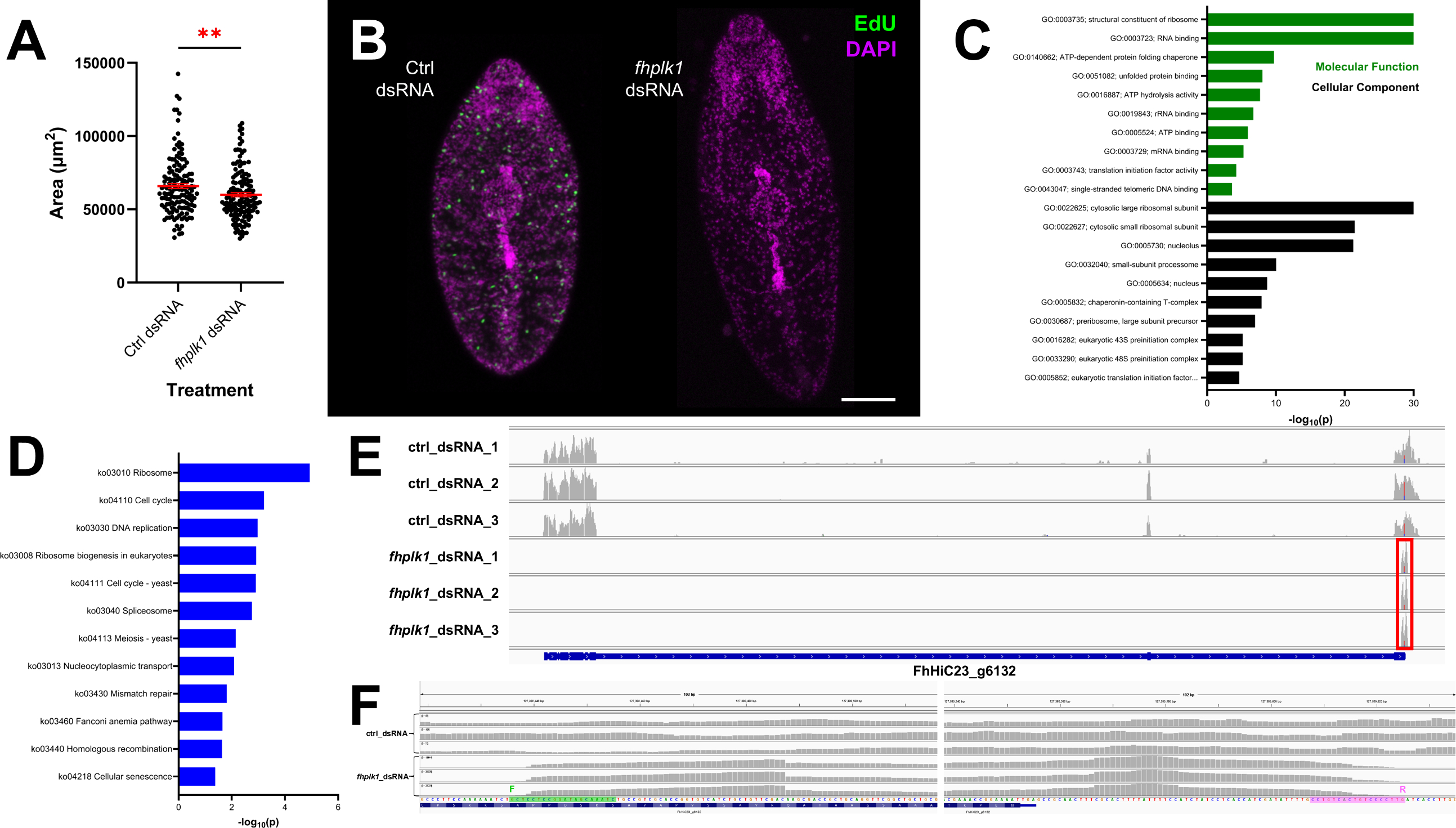

### S4_Fig

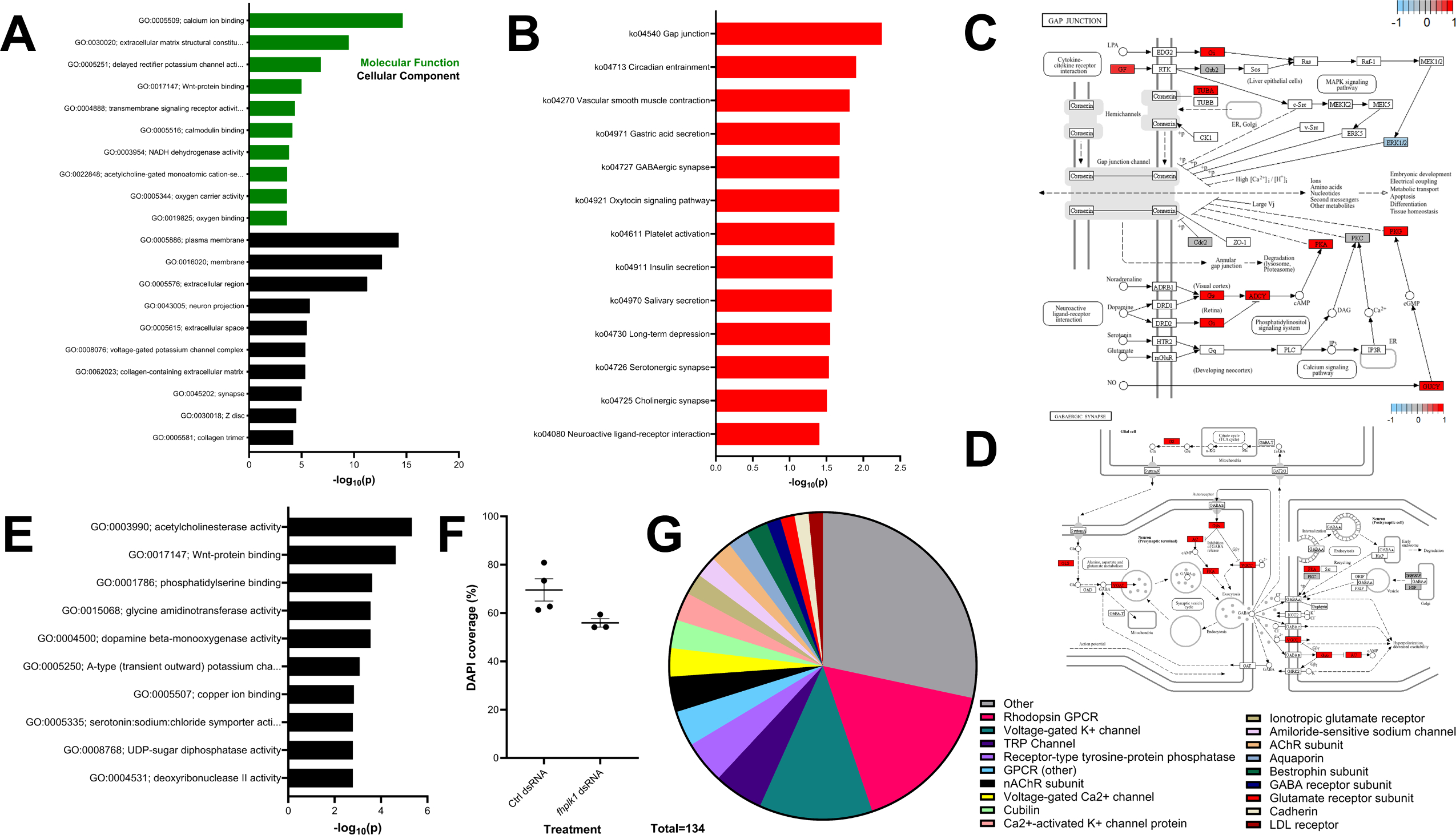

### S5_Fig

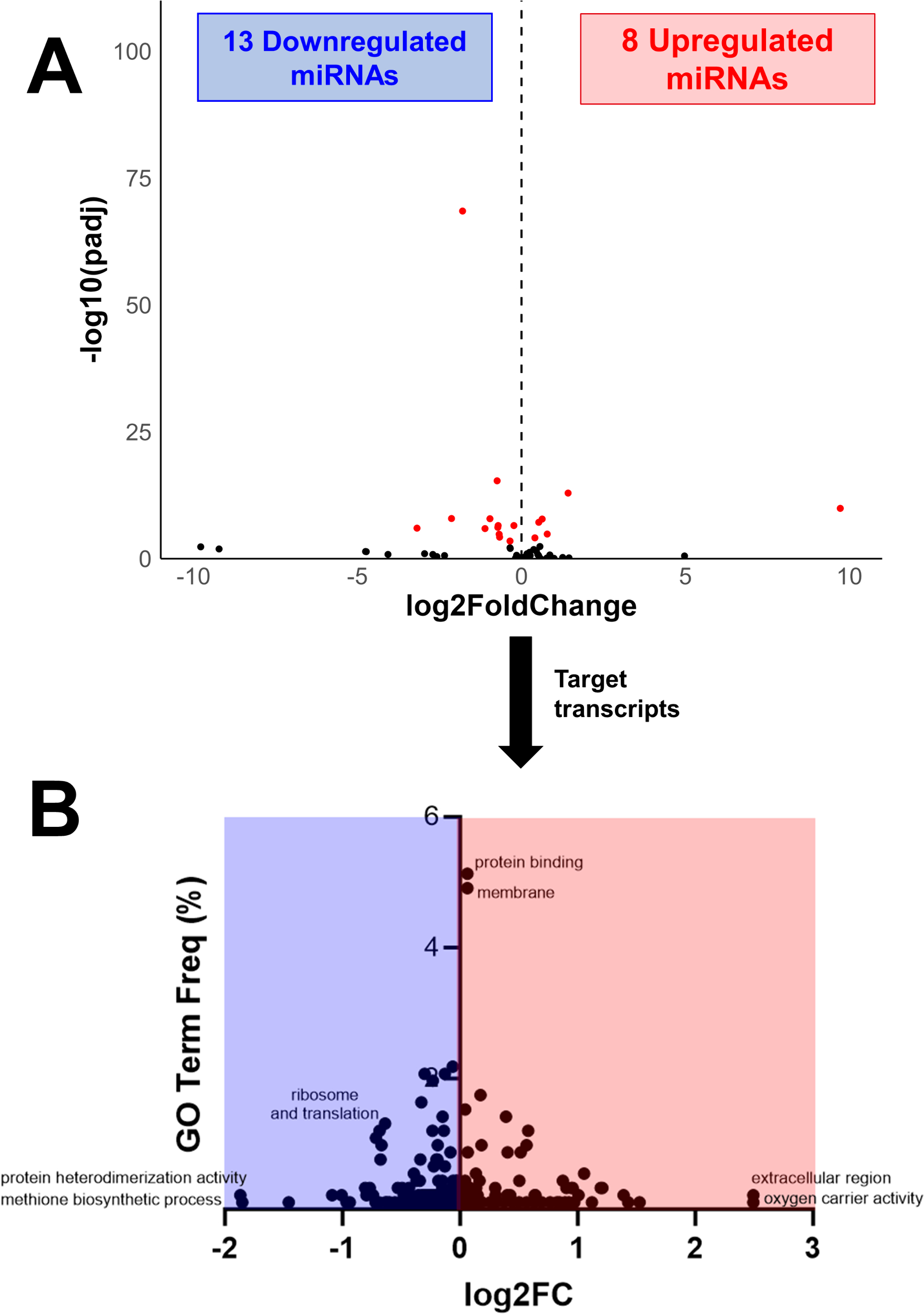

### S6_Fig

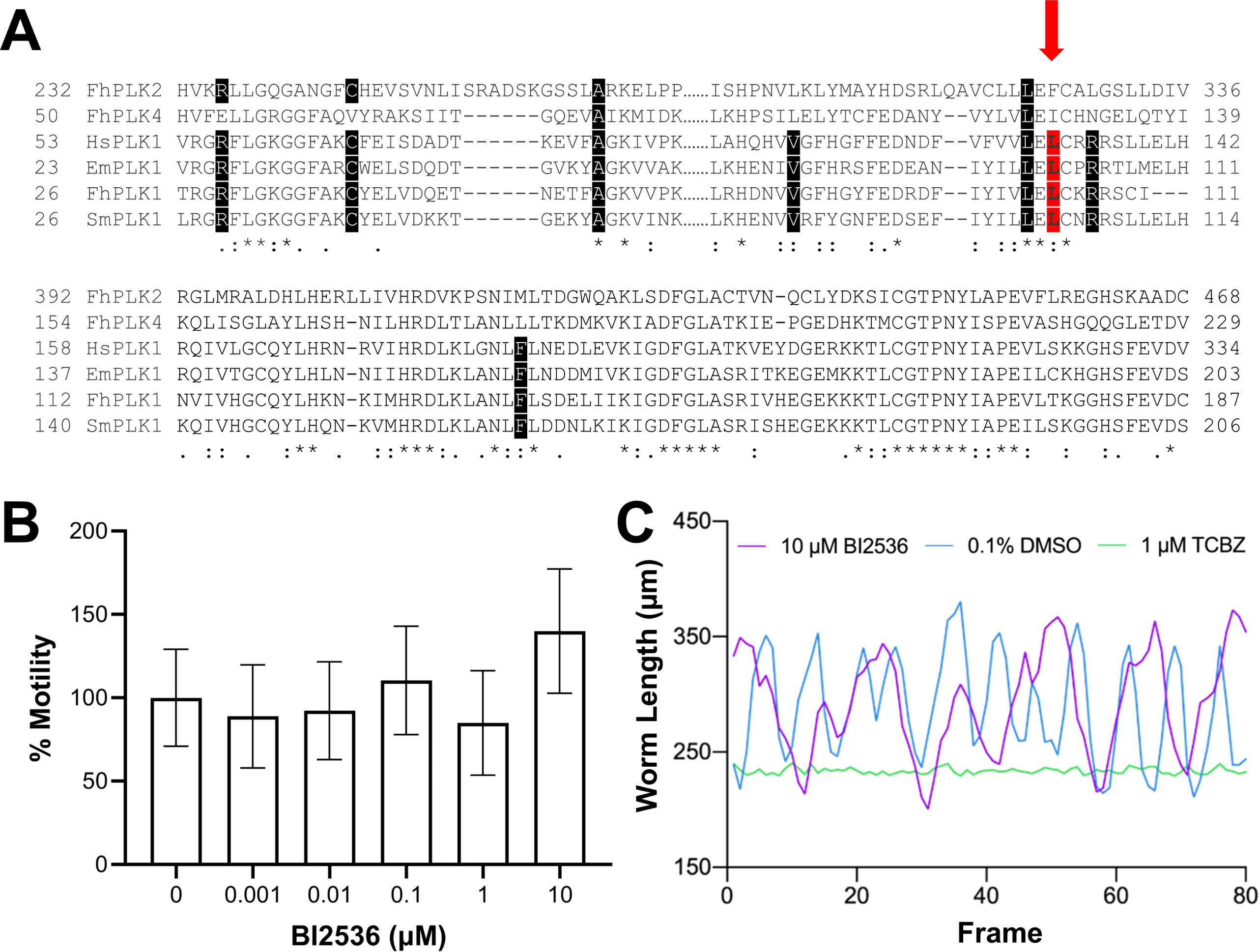
